# Regulation of circular RNA circNFATc3 in cancer cells alters proliferation, migration and oxidative phosphorylation

**DOI:** 10.1101/741041

**Authors:** Thasni Karedath, Fatima M. Al-Dasim, Ikhlak Ahmed, Albandary Al-Qurashi, Afsheen Raza, Simeon S. Andrews, Ayeda Abdulsalam Ahmed, Yasmin Ali Mohamoud, Said Dermime, Joel A. Malek

**Author notes:** These authors contributed equally to this work (Thasni Karedath, Fatima M. Al-Dasim).

## Abstract

Circular RNAs were once considered artifacts of transcriptome sequencing but have recently been identified as functionally relevant in multiple cancers. Although there is still no clear main function of circRNAs, several studies have revealed that circRNAs are expressed in a variety of eukaryotic organisms, demonstrate conservation across species, and are expressed in a regulated manner often independent of their parental linear isoforms. circNFATC3, an abundant and uncharacterized circular isoform of NFATC3 gene (which is formed form backsplicing of exon 2 and 3) in solid tumors was identified from transcriptomic data. Here we show that circNFATC3 gain of and loss of function experiments using RNAi mediated circRNA silencing and circular mini vector-mediated overexpression of circularized constructs in breast and ovarian cancer cell lines affects molecular phenotypes. Knockdown of circNFATC3 induces a reduction in cell proliferation, invasion, migration, and oxidative phosphorylation. Gain of function of circNFATc3 in MDA-MB -231 cells and SKOV3 cells shows a significant increase in cell proliferation, migration, and respiration. The above results suggest that circNFATC3 is a functionally relevant circular RNA in cancer

## Introduction

Circular RNAs (circRNAs) were previously considered transcriptional byproducts; however, they drew attention after the functional characterization of a few circular RNAs *viz*. CDR1as (CiRS-7), SRY, HIPK3 [1][2][3]. CiRS-7, one of the initially identified circRNAs known for its ability to sponge microRNAs, had conceptually changed the understanding of the mechanisms of circRNA and RNA-mediated gene regulation[4]. The wide use of high throughput sequencing analysis has been instrumental in identifying novel circular RNAs in different disease phenotypes and tissues[5][6].

Circular RNAs have attracted increasing attention by cancer researchers as potential biomarkers due to their highly stable nature[7][8][9]. circRNAs have been shown to be differentially regulated in human cancers, including breast, prostate, brain, bladder, colorectal, ovarian, liver and kidney as well as hematological malignancies. Thus, it is becoming increasingly clear that circRNAs have a significant role in cancer pathogenesis, and they are likely to affect several hallmarks of cancer[10,11][12]. Understanding the functional role of circular RNA in cancer cell invasion, migration, tumor suppression provides new insights into cancer detection, oncogenesis, and cancer prevention, which is still elusive. We identified circANKRD12, which is a highly expressed circRNA that regulates tumor invasion and migration in breast and ovarian cancer cells [9]. circNFATc3 was identified by our group as a highly expressed circRNA in metastatic ovarian cancer [8]. The parental gene of circNFATc3 belongs to NFAT gene family which was first identified in immune cells[13–15] and has been associated with malignancies and tumor progression [16,17].

NFATc3 (nuclear factor of activated T cells 3) is a DNA-binding transcription complex consists of a preexisting cytosolic component that translocates to the nucleus upon T cell receptor (TCR) stimulation. NFATc3 plays an important role in retaining stemness *via.* NFATC3/OCT4 signaling and its overexpression increase tumorigenesis in oral cancer [18]. Unlike its parental gene, circNFATc3 remains uncharacterized functionally in cancer. Owing to its abundance in tumor tissues, circNFATc3 might be involved in regulating tumor progression and invasion. Our research focuses mainly on the functional characterization of circNFATc3 in cancer cells by gene silencing and gain-of-function studies.

## Materials and Methods

### Cell Lines and Treatment

Breast cancer cell lines - MDA-MB-231 (ATCC^®^ HTB-26™), MCF7 (ATCC^®^ HTB-22™) - and breast normal cell line - MCF 10A (ATCC^®^ CRL-10317™)-, Primary Mammary Epithelial Cells; Normal, Human (HMEC) (ATCC^®^ PCS-600-010™), ovarian cancer cell lines - PA-1 (ATCC^®^ CRL-1572™), SKOV3 (ATCC^®^ HTB-77™), APOCC (ovarian primary cell line derived from ascites fluid) (pers. communication Dr. Arash Tabrizi), A2780 (93112519-1VL, Sigma), A2780 CIS (93112519-1VL, Sigma), NIH:OVCAR-3 (ATCC^®^ HTB-161™), lung cancer cell line - NCI-H226 (ATCC^®^ CRL-5826™) and lung normal fibroblast cell line LL 24 (ATCC^®^ CCL-151™), T lymphocyte cell Jurkat (ATCC number TIB-152™) (all from American Type Culture Collection, Manassas, VA), were used for the current study. Human lymphoblastoid cell lines (LCL) derived from healthy (LCL-H2), and triple-negative breast cancer patient (LCL-TNBC) were isolated in Dr. Said Dermime’s lab (National Center for Cancer Care and Research, Hamad Medical Corporation, Doha, Qatar). Cells were cultured in DMEM or RPMI (Life Technologies, NY, USA) supplemented with 10% fetal bovine serum (Life Technologies, USA). Low passage number cells were used for all the experiments. Cell culture was routinely checked for mycoplasma contamination using MycoAlert Mycoplasma detection kit (Lonza, Basel, Switzerland). pcDNA3.1 plasmid and pcDNA3.1(+) CircRNA Mini Vector (Addgene) were used to carry out over-expression experiments.

### NFATc3 Gene Expression

NFATc3 circular and linear transcripts were investigated in 15 cell lines - MCF7, MDA-MB-231, SKOV3, LL 24, MCF 10A, NCIH2315, APOCC, AA2780cis, A2780, Ovcar3, PA-1, HMEC, LCL-TNBC, LCL-H2, Jurkat – using beta-actin and B2M as internal controls. RNA isolation, cDNA synthesis, and RT-qPCR were done, as mentioned in the following sections.

### Silencing and Over-expression

Cell viability was tested/checked using Trypan Blue and TC20™ Automated Cell Counter (BIORAD). Cells were seeded at density (5×10^5^) cells/well in 6-well plates. For NFATc3 silencing, siRNA transfection was carried out using two custom-designed siRNAs for both NFATc3 circular and linear transcripts; sicircNFATc3-1, sicircNFATc3-2, silinNFATc3-1, silinNFATc3-2 and universal scrambled control and si-circular junction scrambled controls specific to circular RNA (Supplementary File1). The cells were transfected after 24h with 30pmol concentration of siRNA (Integrated DNA Technologies) or scrambled control (Mission siRNA Universal Negative Control) using Lipofectamine^®^ RNAiMAX Reagent (Invitrogen/Life Technologies) in Gibco™ OptiMEM. For NFATc3 over-expression, cells were transfected with 2.5 micrograms of each empty vectors and vectors containing NFATc3 insert using Lipofectamine^®^ 3000 Transfection Kit (Invitrogen) and treated with G418 for more than two weeks to have a stable transfection.

### Over-expression Vector Preparation

pcDNA3.1(+) CircRNA Mini Vector (plasmid number 60648) [19] and pcDNA3.1 plasmids both obtained from Addgene plasmid repository were used for expressing NFATc3 in cancer cells. circNFATc3 PCR products (Exon 2 and 3) were used for cloning templates. Cloning was conducted according to manufactures protocol using suitable restriction enzymes.

### Nuclear or Cytoplasmic RNA Isolation

After 48h transfection, RNA was isolated using SurePrep™ Nuclear or Cytoplasmic RNA Purification Kit (Fisher Scientific) as described earlier [9].

### RNA Preparation/Isolation and RT-qPCR

Total RNA was isolated from whole cell lysate using miRNeasy Mini Kit (Qiagen) then quantified using Qubit RNA HS Assay Kit (Life technologies). cDNA synthesis was done using random primers for circRNA experiments from iScript™ Select cDNA Synthesis Kit (Bio-Rad). Fast Start Universal SYBR Green Master Mix (Roche) was used to amplify the specific genes using cDNA primers obtained from Integrated DNA Technologies (IDT). Each Real-Time assay was done/run in triplicate using the StepOnePlus Real-Time PCR System (Applied Biosystems) with various primer constructs (IDT).

### RNA-Seq Analysis

After 48h transfection, RNA isolation and DNase digestion using miRNeasy Mini Kit and RNase-Free DNase Set (Qiagen) were done to have/isolate/extract pure RNA. RNA quality control measurement was done using the High Sensitivity RNA Kit and RNA 6000 Nano Kit (Agilent Technologies). Ovation^®^ RNA-Seq System V2 (NuGEN) was used to prepare SPIA cDNA. Libraries were multiplexed using NEXTflex™ DNA Barcodes (Biooscientific) for RNA-Seq. RNA-seq library preparation, In silico detection of circRNA candidates and differentially regulated genes from paired-end RNA-seq data were conducted as described earlier[8][20].

### Cell Proliferation Assay

Cells were seeded at density (5×10^3^) cells/well in flat-bottom 96-well plates. For NFATc3 silencing, cells were transfected after 24h with siRNAs for both NFATc3 circular and linear transcripts then were incubated in 37° CO_2_ injected incubator for (48h, 72h, 96h, and 120h). For NFATc3 over-expression, cells were transfected after 24h with empty vectors and vectors having NFATc3 insert. CellTiter 96^®^ AQueous One Solution Cell Proliferation Assay and CellTiter-Glo 3D Cell Viability Assay (Promega) were conducted according to the manufacturer’s instructions. Luminescence was measured by EnVision Multilabel Plate Reader (PerkinElmer).

### Scratch Assay-Cell Migration Assay

Cells were plated in rectangular cell culture plates using Cell Comb™ Scratch Assay (Merck) and grown to 100% confluency. A wound was created using a cell comb then the medium was replaced with Gibco™ OptiMEM, reduced serum medium, no phenol red. Cells were transfected with respective siRNAs and vectors as mentioned earlier. Cells were fixed with absolute ethanol and stained with crystal violet dye. The distance between the two sides of the cell-free area was photographed using 10X objective AXIO Zeiss epifluorescence microscope. The distance was measured using Zeiss Zen Microscope software (Carl Zeiss Carpenteria, CA, USA).

### Trans-well/Cell Migration and Invasion Assay

Cellular migration and invasion were determined using Corning^®^ Matrigel^®^ Invasion Chamber 6-Well Plate 8.0 Micron (Corning). 10% FBS DMEM was added to the lower chamber as a chemoattractant. Transfected cells were resuspended in serum-free DMEM, transferred to the upper chamber of the Matrigel, incubated for 24h, then visualized under the microscope to count invading cells.

### 3D Model Experiments

3D anchorage-independent spheroids were developed in MDA-MB-231 and HEK293 cell lines by seeding the cells in ultra-low attachment 6-well plates (Corning) for five days to facilitate spheroid formation. Reverse transfection of the spheroids with siRNAs was conducted as described earlier[9].

### Cell Proliferation Assay in 3D Models

Cells were seeded at density (1×10^5^) cells/well in 96-well ultra-low attachment plates with Gibco™ OptiMEM, reduced serum medium, no phenol red. Once spheroids were formed after 5 days, transfection was conducted with siRNAs. After 48h of transfection, CellTiter 96^®^ AQueous One Solution Cell Proliferation Assay was conducted.

### Collagen Invasion Assay in 3D Model

copGFP Control Plasmid:sc-108083 (Santa Cruz Biotechnology) was used to express GFP in the cells. After 48h of transfection, puromycin selection was done for the cells. copGFP transfected MDA-MB-231 cells were seeded in 6-well ultra-low attachment plates for 3D model formation. The cells were silenced using respective siRNAs. 3D structures were embedded in collagen matrix; Gibco™ Collagen I, Rat Protein, Tail (Fisher Scientific) in Falcon™ Chambered Cell Culture Slides (BD Falcon). The invasiveness was analyzed by visualizing the gel under a fluorescence microscope after 72h.

### Assessment of Mitochondrial Function by Seahorse Extracellular Flux Analyzer

The Mitochondrial Oxygen Consumption Rate (OCR) and Extracellular Acidification Rate (ECAR) in MDA-MB-231 cells were assessed by Agilent Seahorse XF Cell Mito Stress Test Kit and Agilent Seahorse XF Cell Energy Phenotype Test Kit (Agilent Technologies) and run in Agilent Seahorse XFe96 Analyzer (Seahorse Bioscience) as per manufacturers protocol. Cells were seeded at density (5×10^3^) cells/well in Seahorse XF Cell Culture Microplate, transfected after 24h and assay were completed 48h after transfection.

### Statistical Analysis

Statistical analysis for all experiments was based on at least three biological replicates and the error bars were drawn with the standard error of means (SEM). The P-value was calculated by using 1-tailed Student’s t-test.

## Results

### Validation of circNFATc3 in cancer cells

Our previous studies reported that circNFATc3 is one of the most abundant circRNAs in cancer cells [8]. We further noted that circNFATc3 derived from NFATc3 gene exon 2 and 3 was abundant across different cancer cell lines compared to normal cells like HMEC, LL 24 and LCL cells (Figure. 1A). Multiple experiments were performed to confirm circNFATc3 expression (Figure. 1). Five different divergent primers were designed to amplify the backsplice exon junction. Each primer pair produced a single distinct band of expected PCR product size, indicating the presence of the circular junction (Figure. 1B). The backsplice junctional sequences were confirmed by Sanger sequencing (Figure. 1C). The divergent primers; with respect to the genomic sequence, only amplified when the cDNA synthesized by random priming was used as a template. While convergent primers only amplify the linear form on DNA samples, the cDNA samples show amplification of both linear and circular forms. This indicates the circular RNAs are a transcriptional splicing product rather than a form present in the genome (Figure. 1D).

**Figure 1.**
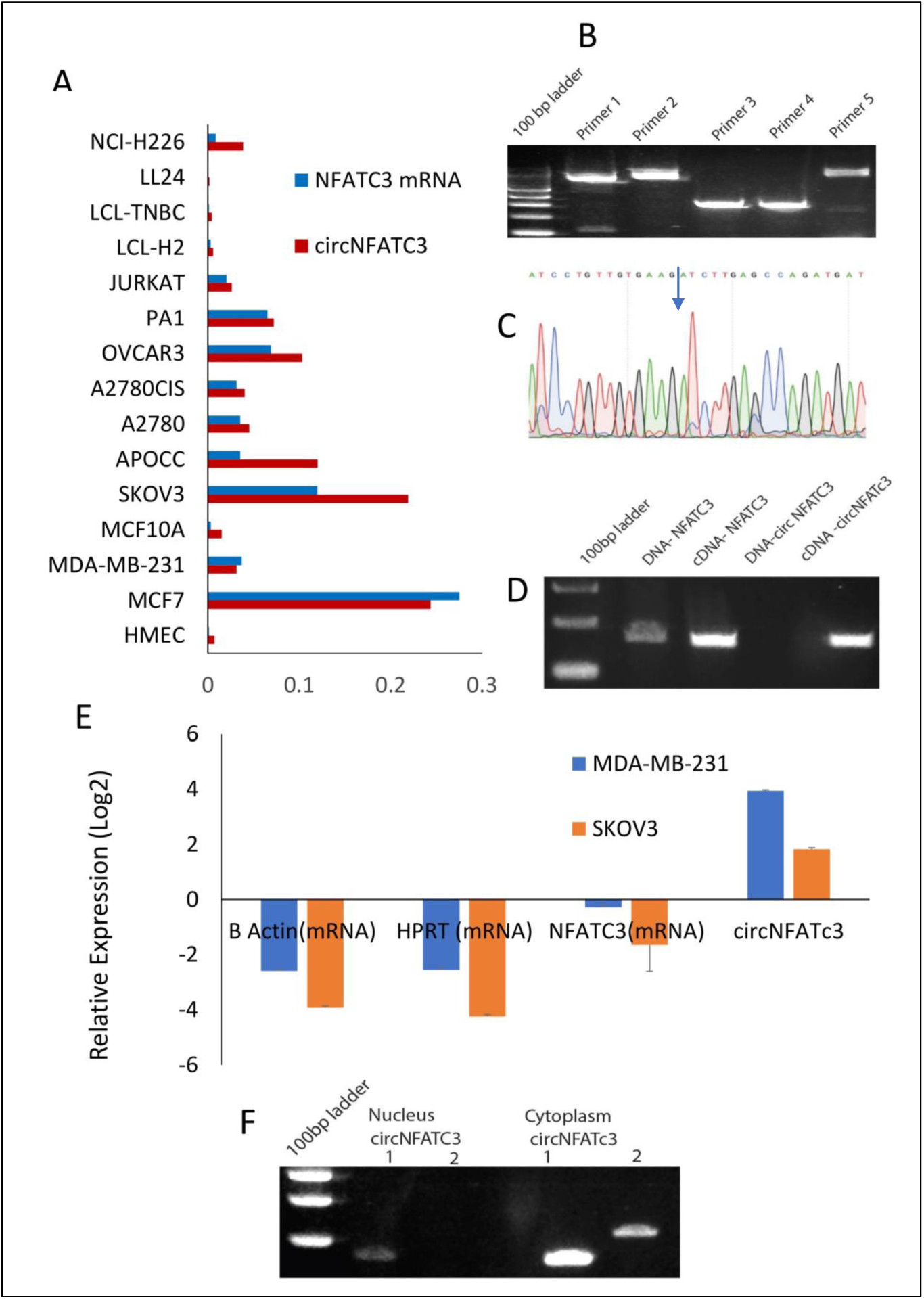
Validation of circNFATc3 expression. **A** Abundance of NFATc3 isoforms in different cell lines. NFATc3 circular and linear transcripts were investigated in 15 cell lines using beta-actin and B2M as internal controls. **B** circNFATc3 backsplice junction amplification. Five different divergent primers were designed to amplify the backsplice exon junction. Each primer pair produced a single distinct band of expected PCR product size indicating the presence of the circular junction. **C** Non-canonical circular splice junction of circNFATc3 exon 3-2. The backsplice junctional sequences were confirmed by Sanger sequencing. **D** Amplification of NFATc3 DNA and cDNA using convergent primers. Convergent primers only amplify DNA samples, the cDNA samples amplifying both indicate the existence of circNFATc3 **E** RNase R resistance of circNFATc3. circRNAs are devoid of 3⍰ single strand overhangs, so they resist RNase R digestion. circNFATc3 is resistant to RNase R digestion compared to linear NFATc3, HPRT, and beta-actin in both cell lines; MDA-MB-231 and SKOV3. **F** Nuclear and cytoplasmic localization of circNFATc3. The PCR amplification of nuclear and cytoplasmic fractions of RNA demonstrated that circNFATc3 is predominantly localized in the cytoplasm.

As circRNAs are devoid of 3⍰ single strand overhangs, they are expected to show resistance to digestion by the exonuclease RNase R. circNFATc3 was resistant to RNase R digestion compared to linear NFATc3, HPRT, and beta-actin in both cell lines. Resistance to digestion with RNase R exonuclease confirms that circNFATc3 is a stable circularized transcript (Figure. 1E). The PCR analysis of nuclear and cytoplasmic fractions of RNA demonstrated that circNFATc3 is predominantly localized in the cytoplasm (Figure. 1F).

We performed Real-Time PCR analysis on 15 different normal and cancer cell lines; including breast, ovarian, lung, lymphoblastoid, to assess the cell-type-specific expression of circNFATc3 (Figure 1A). The majority of cancer cell lines show an abundance of both NFATc3 linear and circular transcripts. Some cancer cells express more circular RNA compared to linear parental mRNA. Breast and ovarian cancer cells show a greater abundance of circNFATc3 compared to normal breast cell lines, lung fibroblast cells, and LCL cells.

### siRNA-mediated silencing of circNFATc3 is highly specific in multiple cancer cells

circNFATc3 originates from Chromosome 16 with the backsplice junction forming between exons 2 and 3 (Figure. 2A). To investigate the functional role of circNFATc3 in cancer cells, we custom designed siRNAs to target the backsplice junction and control siRNAs which have scrambled backsplice junction of circNFATc3 (Figure. 2A, B). These siRNAs were transfected into multiple cancer cell lines to induce siRNA-mediated knockdown of the circular RNA keeping the linear RNA unaffected (Figure. 2C). The circRNA-specific siRNA was designed targeting the backsplice junction spanning exons 2 and 3 of the gene. Three controls were used for the knockdown study, universal scrambled control and circNFATC3 specific scrambled contols as described in the Figure 2B. The circRNA knockdown specificity is demonstrated in Figure 2C and its parental gene knockdown using siRNA downstream of exon3 and exon 2 (in exon 9) left the circRNA unaltered (Figure 2E). MDA-MB-231 cells in an anchorage-independent 3D condition also showed a very high knockdown efficiency (Figure. 2D). We observed high knockdown efficiency ranging between 65% to 95% of the circular junction when using the respective siRNAs versus the controls (scrambled siRNAs) in six cell lines’ transfections (Figure. 2F). Using two siRNA constructs against the circNFATc3, we confirmed that knockdown of the circular RNA is specific and has no significant effect on the linear mRNA expression (Figure. 2B, C). Silencing NFATc3 mRNA using siRNA targeting the linear NFATc3 showed knockdown of NFATc3 mRNA retaining the circNFATc3 intact (Figure. 2E).

**Figure 2.**
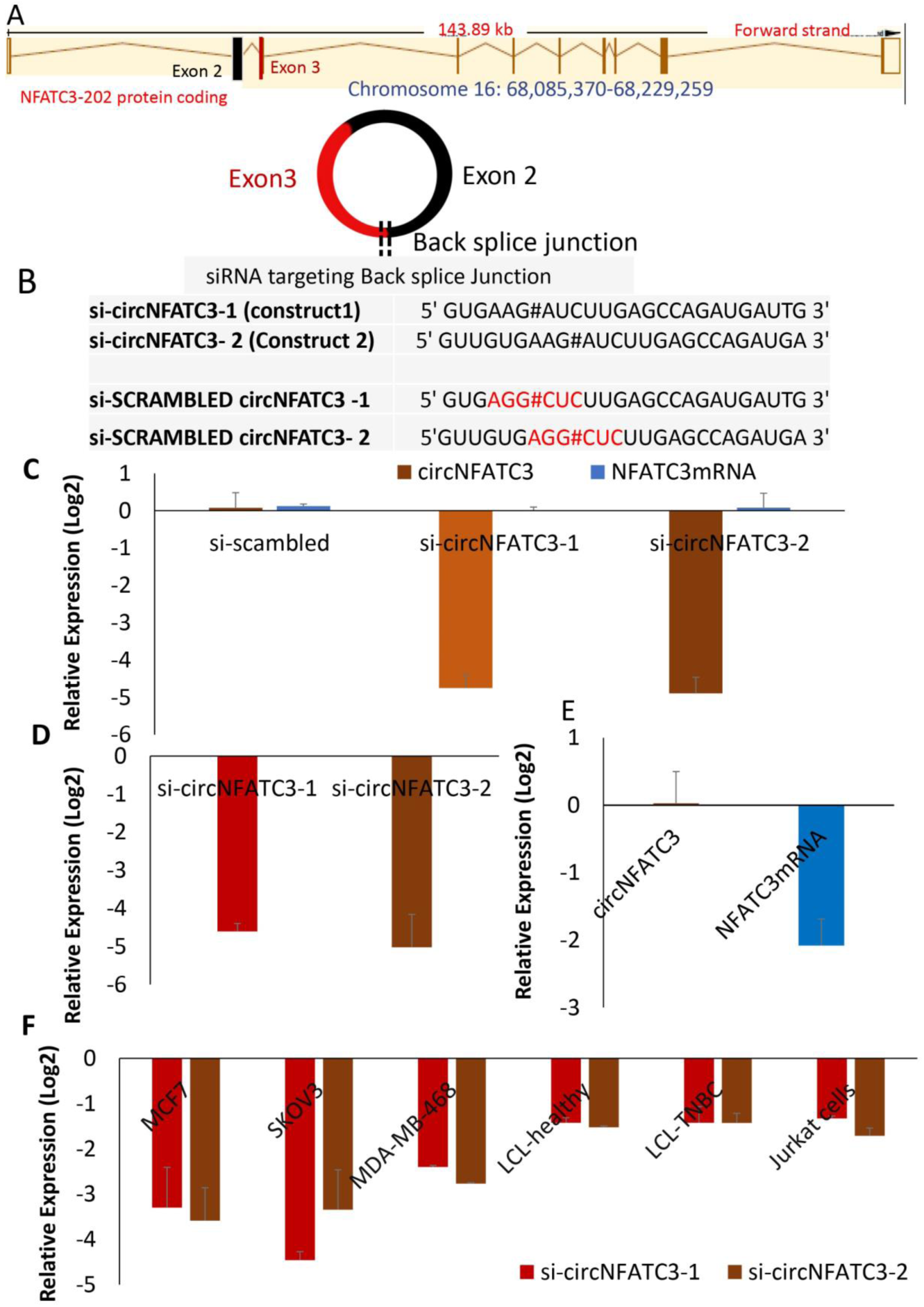
siRNA-mediated functional studies of circNFATc3 in cancer cells. **A** The location of NFATc3 in chromosome 16. The circNFATc3 is formed of exon 2 and 3 by backsplicing. **B** siRNA’ sequences targeting the backsplice junction of circNFATc3 and control siRNAs containing scrambled backsplice junction of circNFATc3. **C** siRNA-mediated knockdown of circNFATc3. siRNAs were transfected into cancer cell line to induce siRNA-mediated knockdown of the circular RNA keeping the linear RNA intact Two different constructs of si-circNFATc3 were used, si-circNFATC3-1,si-circNFATc3-2. **D** Knockdown efficiency of the siRNA-mediated knockdown of circNFATc3 in MDA-MB-231 3D model. **E** siRNA-mediated silencing of NFATc3 mRNA. Silencing NFATc3 mRNA using siRNA targeting the linear NFATc3 preserved/maintained the circNFATc3. **F** siRNA-mediated silencing of circNFATc3 in different cell lines. Six cell lines; that were transfected with two si-circNFATc3 constructs versus scrambled siRNAs showed 65% to 95% knockdown efficiency of the circular junction.

### Silencing of circNFATc3 changes molecular phenotypes of MDA-MB-231 breast cancer cells

RNA-sequencing was performed in three biological replicas of the MDA-MB-231 cell line for both scrambled siRNA control and two siRNA constructs targeting circNFATc3. Number of differentially expressed genes in circNFATc3 knockdown samples with at least 2-fold (log2) change in expression and differentially regulated genes in circNFATc3 silenced MDA_MB 231 in comparison with control cells were listed in Supplementary File 1.

The canonical pathways analysis by Ingenuity IPA toolkit (IPA, QIAGEN Redwood City, (https://www.qiagenbioinformatics.com/products/ingenuity-pathway-analysis/) revealed enrichment of differentially regulated genes involved in cell to cell contact, synthesis of lipids, growth of tumor, cell movement and metabolic pathways (oxidative phosphorylation) (Figure 3,A,B,C,D)(Supplementary Table1). We observed downregulation of EGF, ID1, ID3, and AKT1 which regulate cell to cell contact and cellular movement. Silencing of circNFATc3 affects cellular bioenergetics by downregulating the key genes involved in oxidative phosphorylation and mitochondrial dysfunction (Table 2, Figure 3D). Real-Time validation of RNA-seq data was done for IDH2, ID1, KRT80, and CALCR genes both in circNFATc3 silenced MDA-MB-231 breast cancer cells and SKOV3 ovarian cancer cells (Supplementary Figure1). Transcriptome analysis of differentially expressed genes in circNFATc3 silenced breast cancer cell lines suggested a strong molecular phenotype and we proceeded with the functional screening of circNFATc3 silenced cells using cell-based phenotypic assays in MDA-MB-231 and SKOV3.

**Table 1.** Number of differentially up- and downregulated genes in siRNA-mediated knockdown of circNFATc3 in MDA-MB-231 cells

| Genes downregulated | Genes upregulated |
| --- | --- |
| 839 | 43 |

**Table 2.**
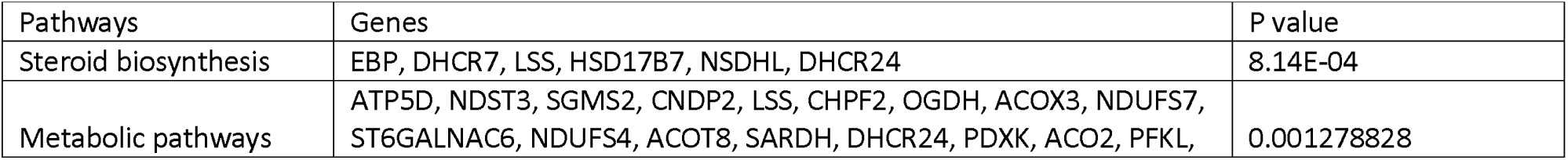
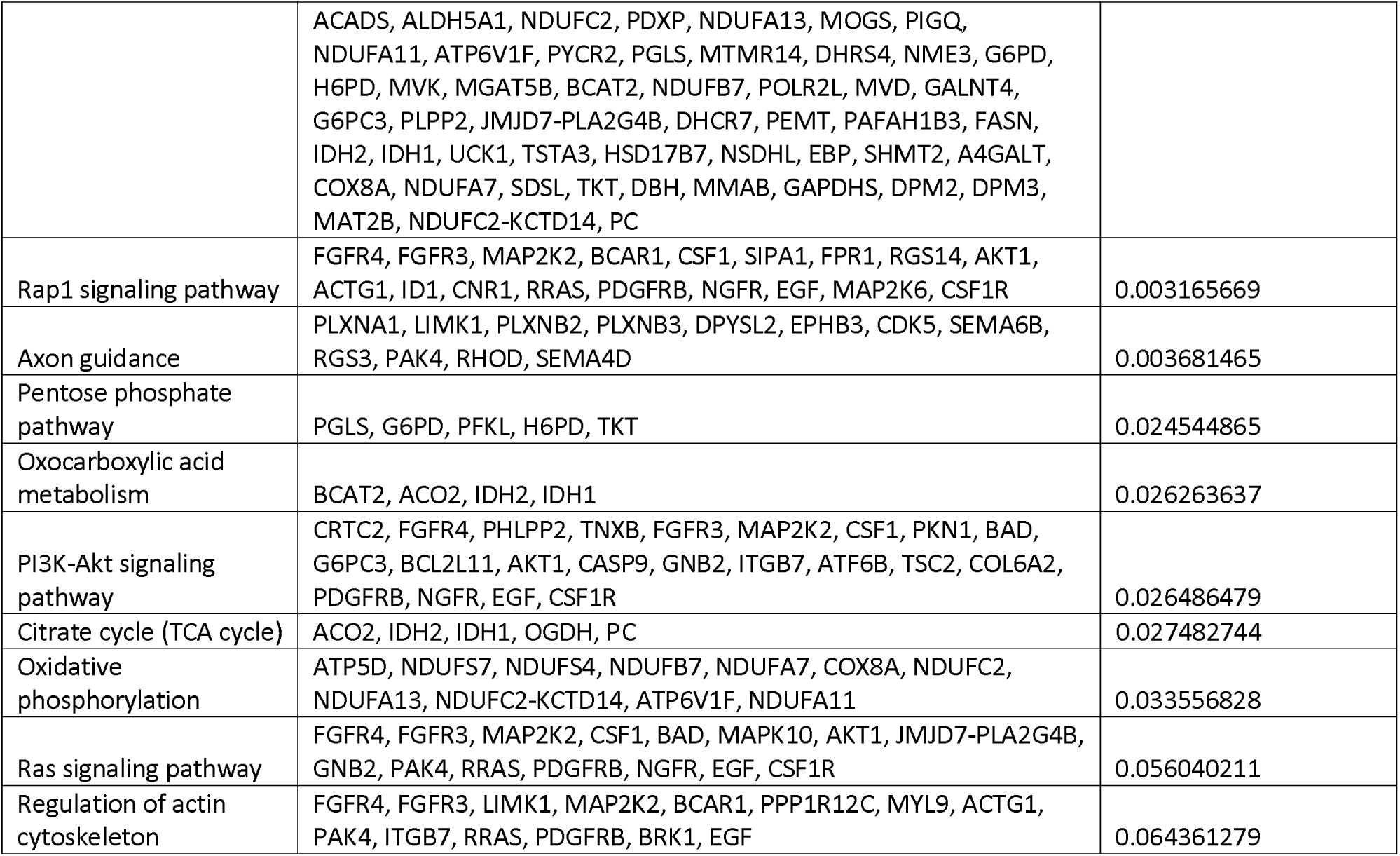
KEGG pathways showing differentially regulated pathways and genes involved in circNFATc3 silenced MDA-MB231 cells

| Pathways | Genes | P value |
| --- | --- | --- |
| Steroid biosynthesis | EBP, DHCR7, LSS, HSD17B7, NSDHL, DHCR24 | 8.14E-04 |
| Metabolic pathways | ATP5D, NDST3, SGMS2, CNDP2, LSS, CHPF2, OGDH, ACOX3, NDUFS7, ST6GALNAC6, NDUFS4, ACOT8, SARDH, DHCR24, PDXK, ACO2, PFKL, | 0.001278828 |
|  | ACADS, ALDH5A1, NDUFC2, PDXP, NDUFA13, MOGS, PIGQ, NDUFA11, ATP6V1F, PYCR2, PGLS, MTMR14, DHRS4, NME3, G6PD, H6PD, MVK, MGAT5B, BCAT2, NDUFB7, POLR2L, MVD, GALNT4, G6PC3, PLPP2, JMJD7-PLA2G4B, DHCR7, PEMT, PAFAH1B3, FASN, IDH2, IDH1, UCK1, TSTA3, HSD17B7, NSDHL, EBP, SHMT2, A4GALT, COX8A, NDUFA7, SDSL, TKT, DBH, MMAB, GAPDHS, DPM2, DPM3, MAT2B, NDUFC2-KCTD14, PC |  |
| Rap1 signaling pathway | FGFR4, FGFR3, MAP2K2, BCAR1, CSF1, SIPA1, FPR1, RGS14, AKT1, ACTG1, ID1, CNR1, RRAS, PDGFRB, NGFR, EGF, MAP2K6, CSF1R | 0.003165669 |
| Axon guidance | PLXNA1, LIMK1, PLXNB2, PLXNB3, DPYSL2, EPHB3, CDK5, SEMA6B, RGS3, PAK4, RHOD, SEMA4D | 0.003681465 |
| Pentose phosphate pathway | PGLS, G6PD, PFKL, H6PD, TKT | 0.024544865 |
| Oxocarboxylic acid metabolism | BCAT2, ACO2, IDH2, IDH1 | 0.026263637 |
| PI3K-Akt signaling pathway | CRTC2, FGFR4, PHLPP2, TNXB, FGFR3, MAP2K2, CSF1, PKN1, BAD, G6PC3, BCL2L11, AKT1, CASP9, GNB2, ITGB7, ATF6B, TSC2, COL6A2, PDGFRB, NGFR, EGF, CSF1R | 0.026486479 |
| Citrate cycle (TCA cycle) | ACO2, IDH2, IDH1, OGDH, PC | 0.027482744 |
| Oxidative phosphorylation | ATP5D, NDUFS7, NDUFS4, NDUFB7, NDUFA7, COX8A, NDUFC2, NDUFA13, NDUFC2-KCTD14, ATP6V1F, NDUFA11 | 0.033556828 |
| Ras signaling pathway | FGFR4, FGFR3, MAP2K2, CSF1, BAD, MAPK10, AKT1, JMJD7-PLA2G4B, GNB2, PAK4, RRAS, PDGFRB, NGFR, EGF, CSF1R | 0.056040211 |
| Regulation of actin cytoskeleton | FGFR4, FGFR3, LIMK1, MAP2K2, BCAR1, PPP1R12C, MYL9, ACTG1, PAK4, ITGB7, RRAS, PDGFRB, BRK1, EGF | 0.064361279 |

**Figure 3.**
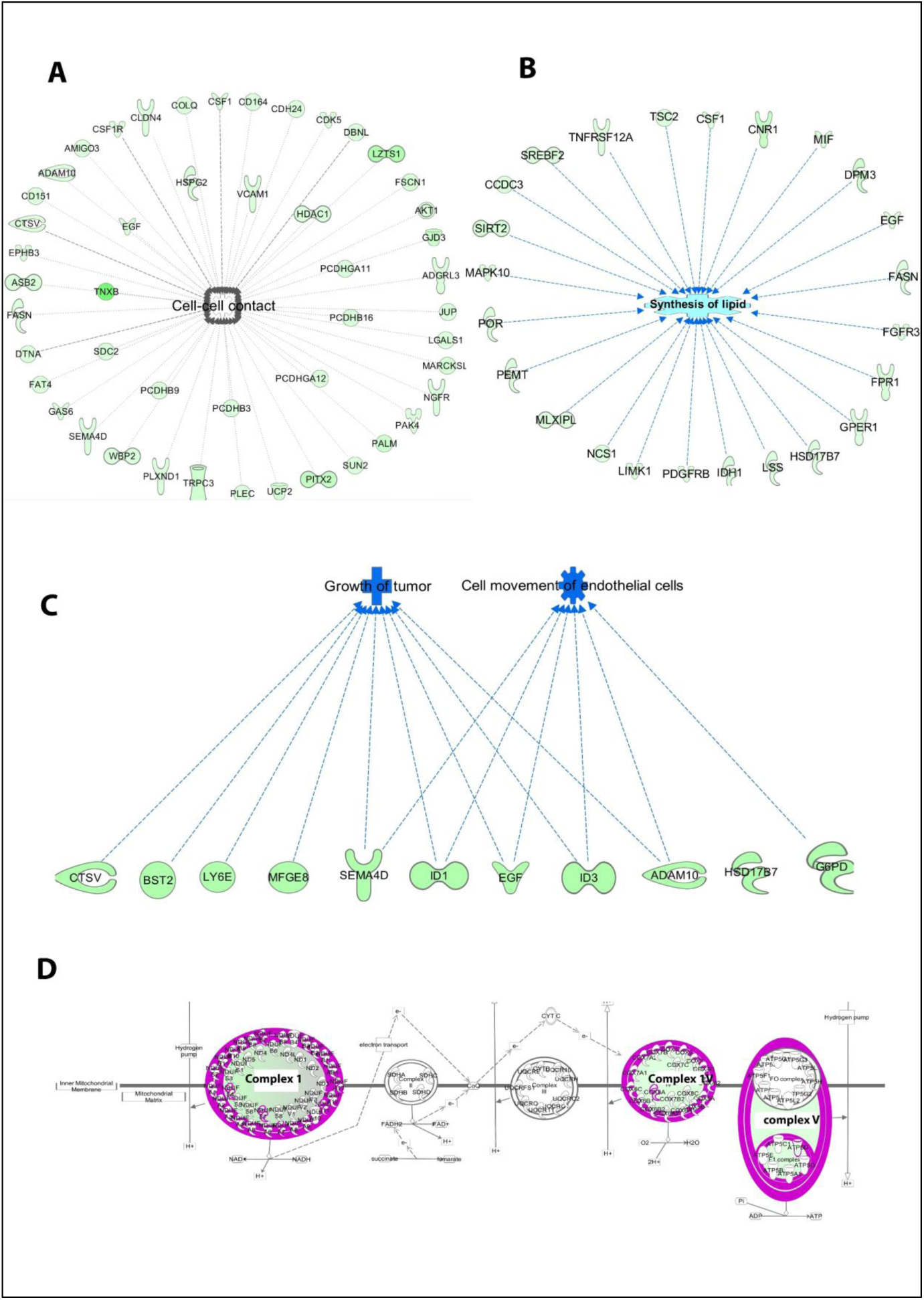
RNA-seq analysis-differentially expressed genes in circNFATc3 silenced MDA-MB-231 cells (circNFATc3 vs control). The canonical pathways analysis by Ingenuity^®^ IPA toolkit identified enrichment of differentially regulated genes involved in many pathways including: **A** Cell-cell contact. **B** Lipid synthesis. **C** Growth of tumor and cell movement. **D** Oxidative phosphorylation/mitochondrial dysfunction.

### siRNA-mediated silencing of circNFATc3 decreases cell proliferation, migration and invasion in breast and ovarian cancer cells

CircNFATc3 silenced cells show a significant reduction in cell proliferation compared to cells with the scrambled siRNA controls in MTS cell proliferation and ATP assays (Figure 4). This reduction was seen in both MDA-MB-231 and SKOV3 cells (Figure 4A, B). The cell proliferation assays viz; MTS and ATP, indicating that the silencing of the NFATc3 parental gene can induce a significant reduction in cell viability but not as substantial as its circular counterpart. Both cell lines show a significant reduction in cell proliferation after 72h of transfection, however MDA-MB-231 cells were able to show a reduction in cell proliferation as early as 48h (Figure 4A) (These results suggest that knockdown of circular forms of NFATc3 gene is capable of inducing strong phenotypic changes and modulate the growth of cancer cells.

**Figure 4.**
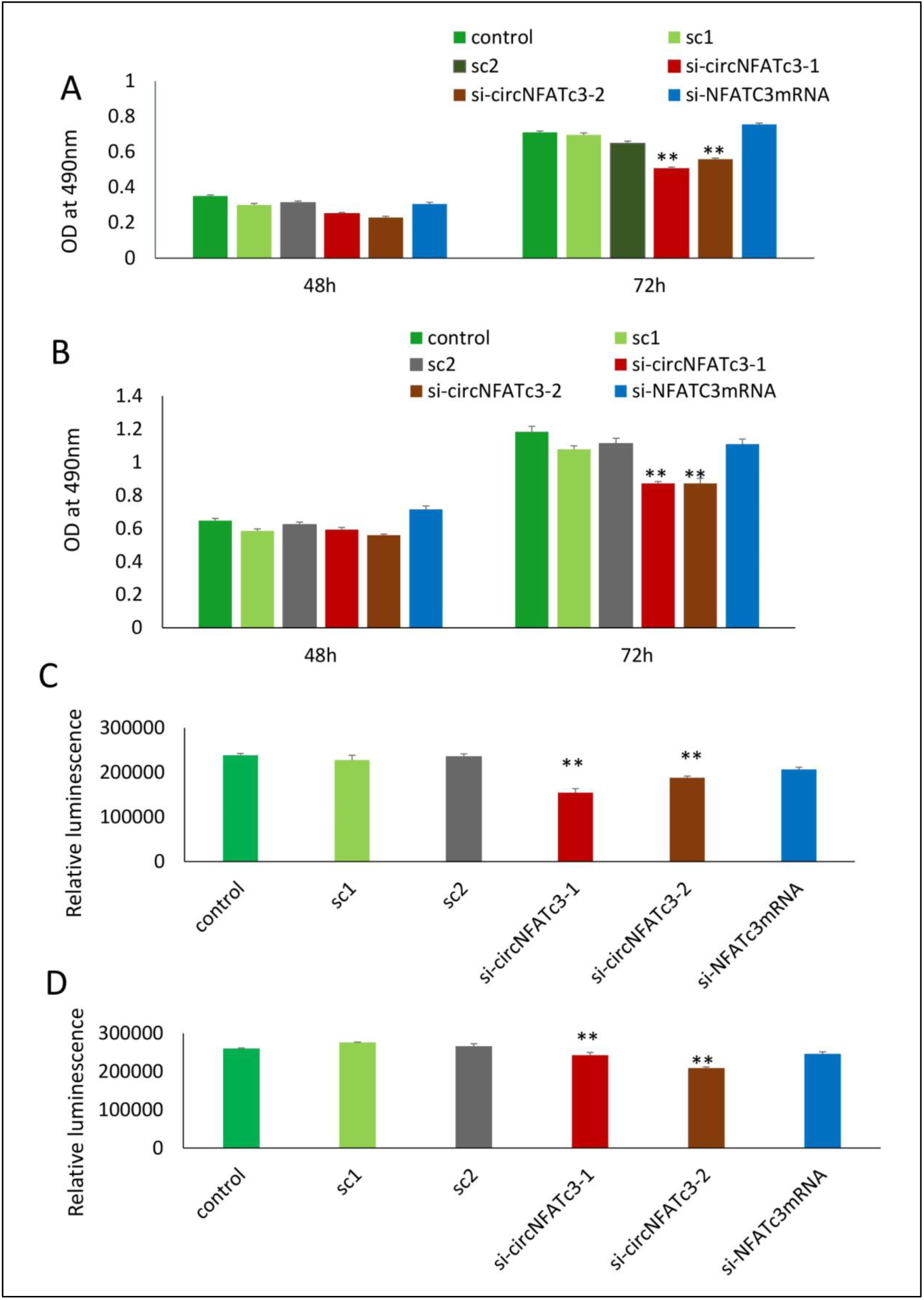
Cell proliferation (MTS and ATP) assays in circNFATc3 and NFATc3 mRNA silenced cells. The cell proliferation assays indicate that silencing of NFATc3 parental gene can induce a significant reduction in cell viability but not as substantial as its circular counterpart. **A** MTS assay of NFATc3 silenced MDA-MB-231 cells. **B** MTS assay of NFATc3 silenced SKOV3. **C** ATP assay of NFATc3 silenced MDA-MB-231 cells. **D** ATP assay of NFATc3 silenced SKOV3 cells. Data in **B** and **D** are the means with error bars indicating standard error of the mean (SEM) of three experiments/biological replicates. \*\**P*⍰<⍰0.01 (Student’s t-test).

Wound healing assay shows that silencing of circNFATc3 reduces cell migration in both MDA-MB-231 and SKOV3 cells (Figure 5A, B), after 48⍰h and 72⍰h compared to the scrambled control. Matrigel invasion (inserts coated with matrigel) analysis using a Boyden chamber shows that circNFATc3 silenced cells undergo a significant reduction in the invasion and migration compared to scrambled control (Figure 5C, D). Collagen invasion assay of the circNFATc3 silenced cells in 3D anchorage-independent condition exhibits less invasion and failed dispersion through collagen matrix compared to scrambled control (Figure 5E).

**Figure 5.**
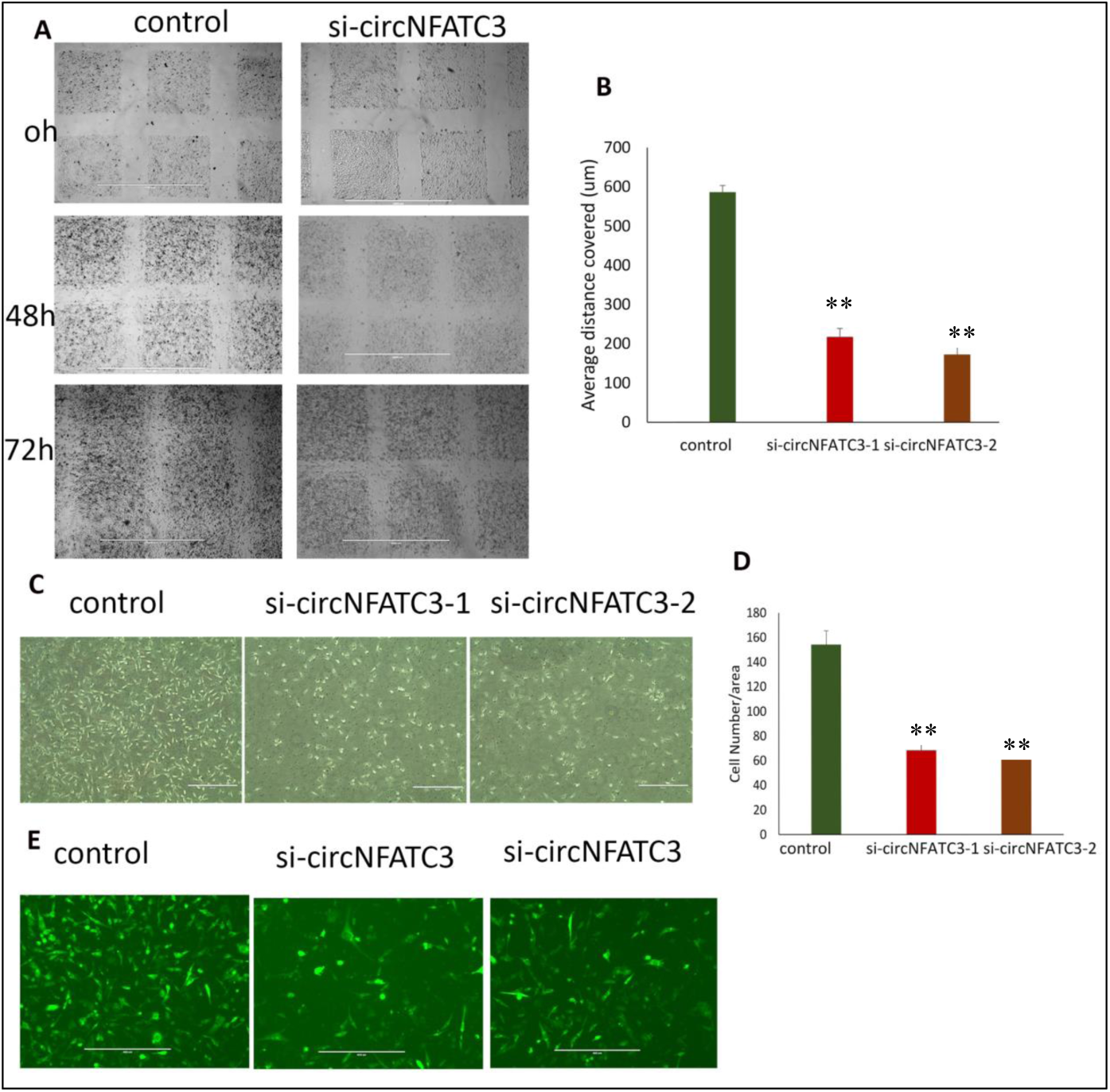
Migration and invasion assays of circNFATc3 silenced MDA-MB-231 cells. A Wound healing assay. circNFATc3 silenced cells reduce migration after 48h and 72h compared to control. B Scratch distance covered average. Wound healing analysis shows a significant reduction in migration of circNFATc3 silenced cells compared to control cells. C Boyden chamber. It shows a significant reduction in the invasion and migration of circNFATc3 silenced cells compared to control cells. D circNFATc3 silenced MDA-MB-231 cells and control cells penetration through matrigel; collagen matrix. circNFATc3 silenced cells number that traveled in the denuded space is lesser than the number of the control cells. E Collagen invasion assay of circNFATc3 silenced cells in the 3D model. The circNFATc3 silenced cells exhibit less invasion fail dispersion through the collagen matrix compared to control cells. Data in B and D are the means with error bars indicating standard error of the mean (SEM) of three experiments. **P⍰<⍰0.01 (Student’s t-test).

### siRNA-mediated silencing of circNFATc3 modulating cellular bioenergetics shows a shift in metabolic phenotype

RNA-seq analysis revealed that knockdown of circNFATc3 can regulate oxidative phosphorylation and TCA cycle thereby affecting the mitochondrial function directly. We used extracellular flux assays which allow direct evaluation of cellular bioenergetic profiles *ex vivo* by measuring oxygen consumption rate (OCR, a measure of oxidative phosphorylation) and extracellular acidification rate (ECAR) and cell energy phenotype. To functionally validate the RNA-seq results, we performed extracellular flux analysis for mitochondrial potential and cell energy phenotype in MDA-MB-231 and SKOV3 cells (Figure 6A, Supplementary files). As per the RNA-seq data we found that both the respiratory capacity and the aerobic glycolysis, as measured by OCR and ECAR respectively, of circNFATc3 silenced cells were significantly low compared to the scrambled control (Figure 6A, B, C, D, E). This result indicates that circNFATc3 knocked down cells exist in a relatively low bioenergetic state while scrambled control cells adapt an energetic - high respiratory capacity, high glycolysis - metabolic phenotype. (Figure 6A). SKOV3 cells also follow the same trend, as silencing circNFATc3 lowers mitochondrial respiration (Supplementary Figure 2). Taken together, these data show that knocking down circNFATc3 in cancer cells maintains a quiescent metabolic phenotype of low respiratory capacity and glycolysis compared to control cells.

**Figure 6.**
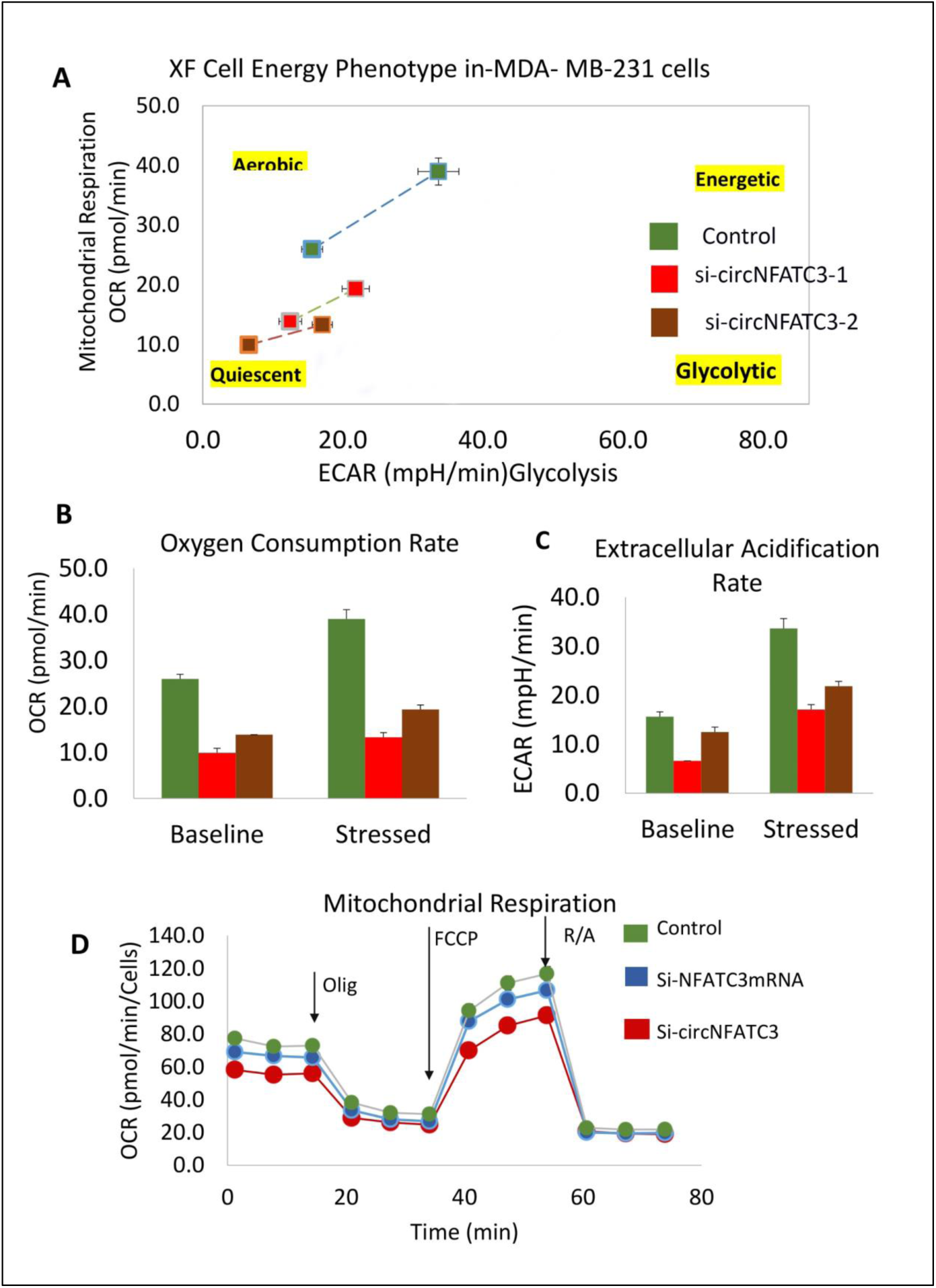
Metabolic alteration in circNFATc3 and NFATc3 mRNA silenced MDA-MB-231 cells. **A** XF cell energy phenotype in circNFATc3 silenced cells. circNFATc3 silenced cells maintain a quiescent metabolic phenotype with low OCR and ECAR compared to control cells. **B** Oxygen consumption rate in circNFATc3 silenced cells. OCR is significantly low in circNFATc3 silenced cells compared to control cells. **C** Extracellular acidification rate in circNFATc3 silenced cells. ECAR is significantly low in circNFATc3 silenced cells compared to control cells. **D** Mitochondrial respiration of the circNFATc3 silenced cells.

### Over-expression of circNFATc3 regulates cell proliferation, migration, and cellular bioenergetics

MDA-MB-231 and SKOV3 cells were used for gain-of-function assays due to their moderately low expression of circNFATc3 and high level of transfection efficiency. The circNFATc3 construct in the pcDNA3.1(+) CircRNA Mini Vector containing ALU repeats[19] can potentially circularize the NFTAc3 construct spanning from Exon 2 to Exon 3. This is in contrast to its parental pcDNA3.1 Vector without the ALU repeats which does not circularize exons. Thus, both vectors were used for ectopic expression of exons 2 and 3 of NFATc3 along with their respective empty vectors in MDA-MB-231 and SKOV3 cells. The NFATc3 linear construct (exon 2 and exon3) in pcDNA3.1 vector, its empty vector, and empty pcDNA3.1(+) CircRNA Mini Vector were used in transfection as controls to identify the phenotypic effect of circNFATc3 in the overexpressed cells. The pcDNA3.1(+) CircRNA Mini Vector with ALU repeats was able to circularize the NFATc3 construct in MDA-MB-231 and SKOV3 cells. MDA-MB-231 cells showed high transfection efficiency and were able to overexpress 16-fold higher levels of circularized NFATc3 construct compared to empty vector with alu repeats(circRNA mini Vector). Thus, we ectopically overexpressed circNFATc3 in MDA-MB-231 and SKOV3 cells using pcDNA3.1(+) CircRNA Mini Vector. For comparison, we also overexpressed the linear form of exon 2 and 3 in the pcDNA3.1 Vector that does not allow subsequent circularization of the structure (Figure. 7A, B). The NFATc3 (exon 2 and 3) construct is 1298 bp long. Only the NFATc3 construct in pcDNA3.1(+) CircRNA Mini Vector with ALU repeats was able to circularize the majority of the transcript (Figure 7A, B).

**Figure 7.**
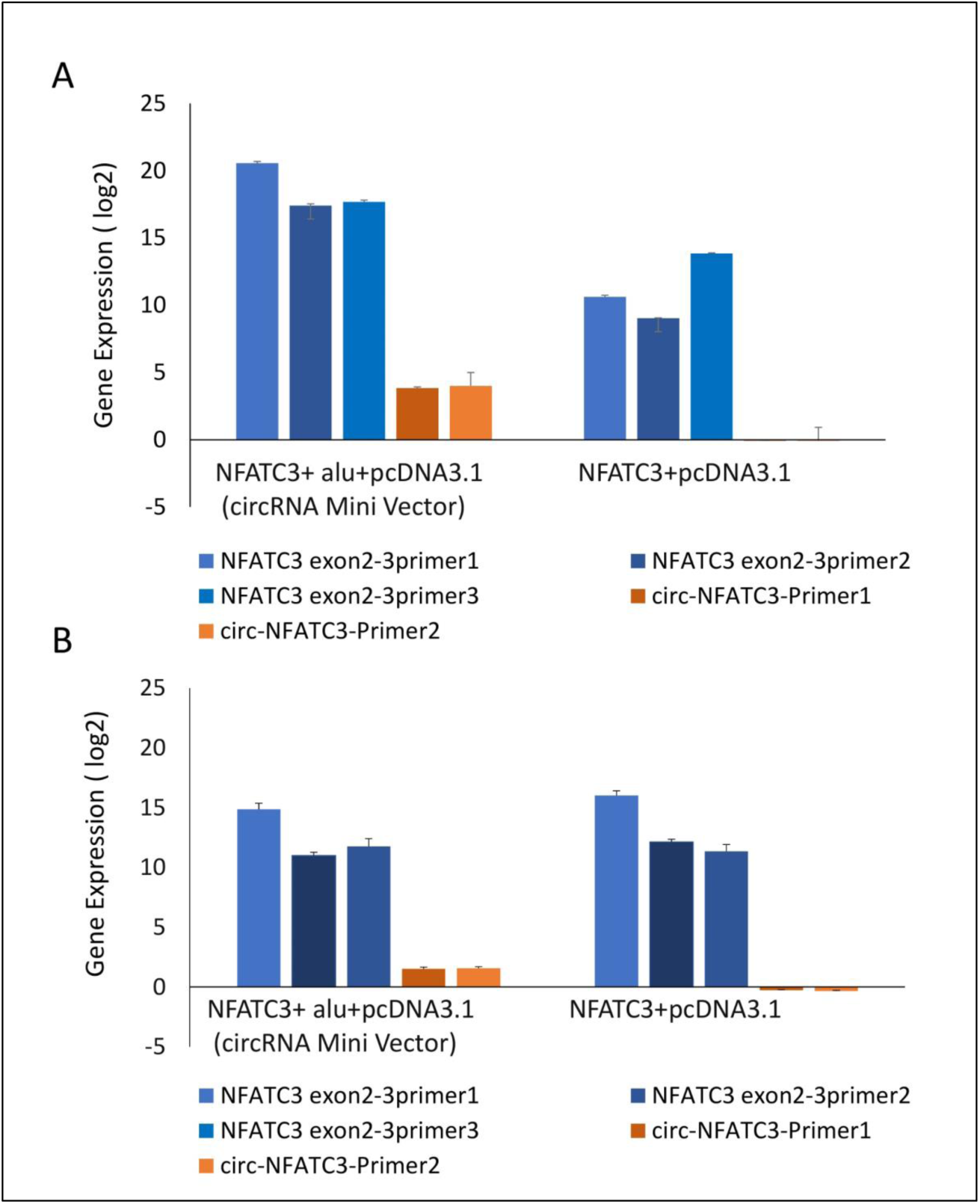
Gain-of-function assay of the NFATc3 gene using pcDNA3.1(+) CircRNA Mini Vector and pcDNA3.1 Vector. NFATc3 construct in the pcDNA3.1(+) CircRNA Mini Vector expressed both circNFATc3 and NFATc3 mRNA, but the pcDNA3.1 Vector was not able to circularize the NFATc3 construct. **A** NFATc3 overexpression in MDA-MB-231 cells. **B** NFATc3 overexpression in SKOV3 cells.

Our next aim was to identify the phenotypic effect of the circularized transcript compared to the non-circularized control transcript. Cell proliferation assays at 48h and 72h revealed increases in both cell lines in circNFATc3 overexpressed groups compared to controls (Figure. 8A, B, C, D). Moreover, overexpression of circNFATc3 dramatically enhanced cell migration in MDA-MB-231 cells (Figure. 8E, F). We further determined the alteration of cellular bioenergetics of the overexpressed circNFATc3 and control in MDA-MB-231 and SKOV3 cells by using extracellular flux analysis (Figure 9, Supplementary Figure 3, 4). As shown in figure 9 a significant increase was observed in OCR and ECAR with a slight shift in energy phenotype only in circNFATc3 overexpressed cells compared to all other control conditions (Figure 9 A,B,C). Taken together, the overexpression of circNFATc3 can increase cancer cell proliferation, migration, and bioenergetics.

**Figure 8.**
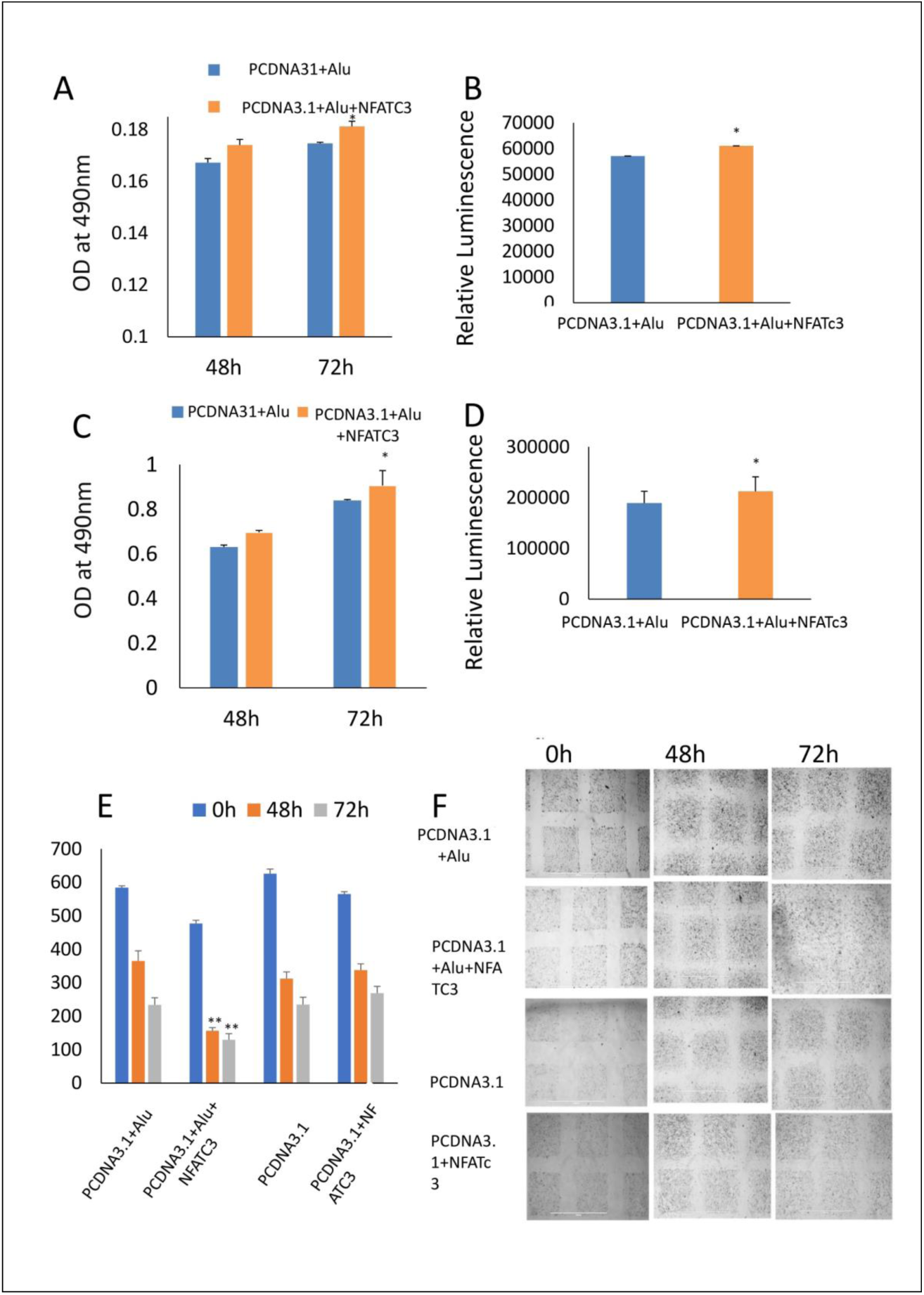
Phenotypic assays of the NFATc3 overexpressed cells. A MTS assay of the NFATc3 overexpressed MDA-MB-231 cells. B ATP assay of the NFATc3 overexpressed MDA-MB-231 cells. C MTS assay of the NFATc3 overexpressed SKOV3 cells. D ATP assay of the NFATc3 overexpressed SKOV3 cells. E Scratch covered distance average measured at 0h, 48h, and 72h. Wound healing analysis shows a significant increase in migration of the circNFATc3 overexpressed cells compared to control cells. F Wound healing assay. circNFATc3 overexpressed cells increase migration after 48h and 72h compared to control. Data in A - E are the means with error bars indicating standard error of the mean (SEM) of three experiments. **P⍰<⍰0.01 (Student’s t-test).

**Figure 9.**
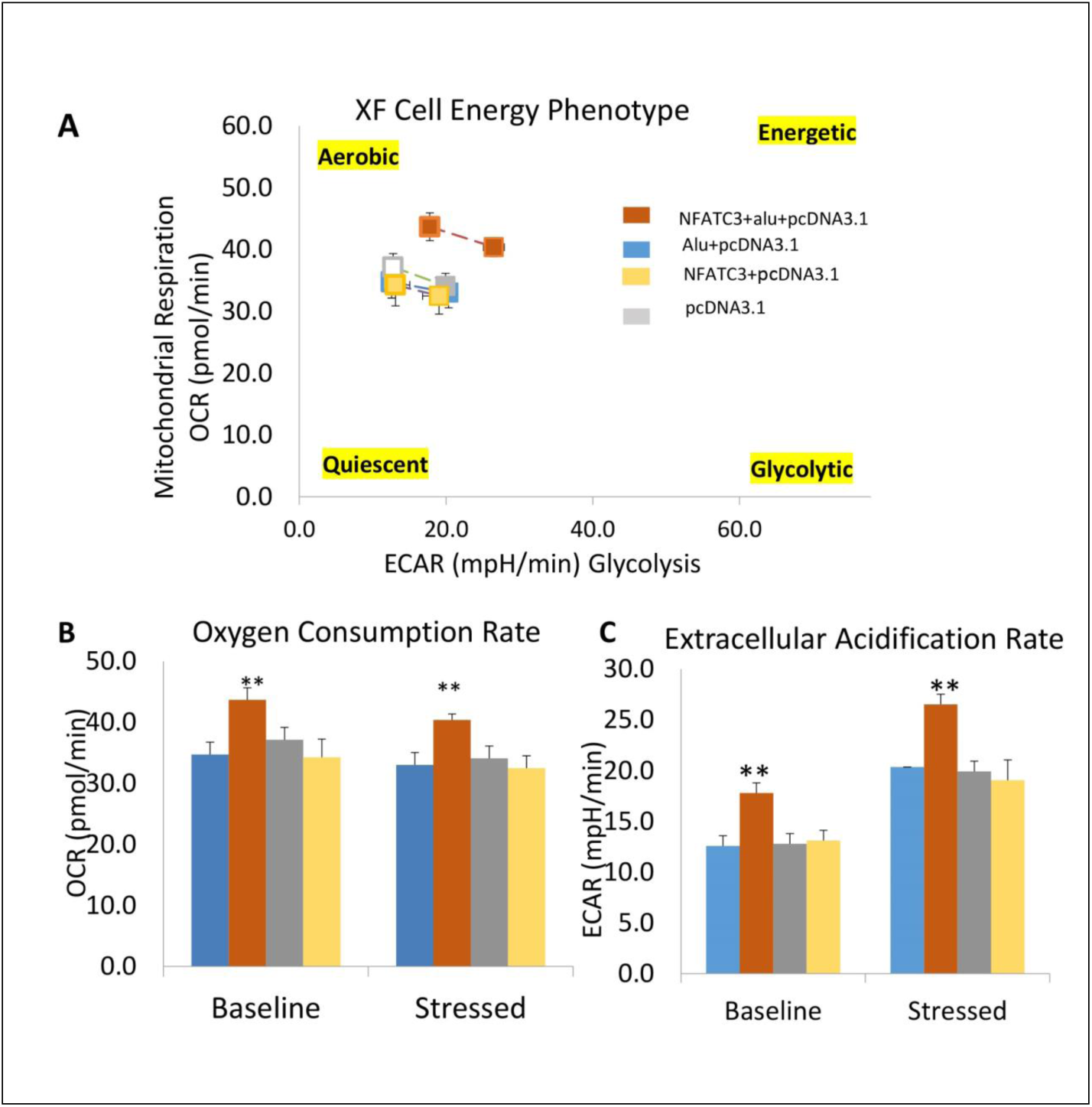
Metabolic alteration in circNFATc3 and NFATc3 mRNA overexpressed MDA-MB-231 cells. A XF cell energy phenotype in circNFATc3 overexpressed cells. circNFATc3 overexpressed cells maintain an aerobic metabolic phenotype with high OCR and ECAR compared to control cells. B Oxygen consumption rate in circNFATc3 overexpressed cells. OCR is high in circNFATc3 overexpressed cells compared to control cells. C Extracellular acidification rate in circNFATc3 overexpressed cells. ECAR is significantly=high in circNFATc3 overexpressed cells compared to control cellsData in B and C are the means with error bars indicating standard error of the mean (SEM) of three experiments. **P⍰<⍰0.01 (Student’s t-test).

## Discussion

Due to the extensive use of high-throughput sequencing platforms to identify novel regulatory RNAs, increasing numbers of circRNAs have been identified in human samples. Emerging evidence demonstrating that circRNAs play crucial roles in carcinogenesis and cancer progression led to rapid exploration of the functional relevance of these RNAs. Similar to oncogenes, aberrantly expressed circRNAs have been reported in diverse cancer types [21–23]. We identified circNFATc3, which is a highly expressed circRNA in breast and ovarian cancer cells [8]. circNFATc3 is characterized and identified as circRNA consisting of two exons with a length of 1298bp and is resistant to RNase R treatment (Figure 1E). The annotated circNFATc3 isoforms listed in circnet (http://circnet.mbc.nctu.edu.tw/.)[24] share the same backsplice junction studied here is highly expressed in different tissues; particularly in breast cancer tissue compared to normal tissue (Supplementary Figure 5). circNFATc3 is annotated as hsa_circ_0000711 (circBase, http://www.circbase.org/cgi-bin/webBlat), NFATc3_hsa-circRNA3069 (starBase, http://starbase.sysu.edu.cn/index.php). Even though circNFATc3 is annotated by different circRNA based databases, it remains functionally uncharacterized. The only documented functional characterization of circNFATc3 is its association with RNA binding protein - IMP3 - potentially involved in the biogenesis of circular RNAs[25].

As a step towards functionally characterizing the circNFATc3 of exon2,3 we conducted RNA silencing without altering the expression of its parental mRNA. Series of experiments were conducted to validate the knockdown efficacy in different cell lines using two siRNA constructs, a universal scrambled control, and two scrambled constructs of circNFATc3 (figure2C). Knockdown of circNFATc3 in MDA-MB-231 and SKOV3 cells that have a moderately high level of circNFATc3 shows a reduction in cell proliferation. However, knockdown of circNFATc3 in LCL (lymphoblastoid) cells which have a low level of circNFATc3 expression was unable to induce any phenotypic changes (supplementary Figure 6, Figure 1A). These results suggest that circNFATc3 knockdown was highly specific to circular RNA, and it has no off-target effect. Transcriptome analysis of circNFATc3 silenced MDA-MB-231 cells compared to scrambled cells shows a distinct molecular phenotype (figure 3). Cell proliferation, cell to cell contact, cell movement and oxidative phosphorylation were altered in circNFATc3 silenced MDA-MB-231 cells (Figure 3). Cell proliferation assays revealed that knockdown of circNFATc3 can significantly reduce cell proliferation, cell migration and cancer invasion (figure 4,5). circRNA junction (backsplice junction) is essential for altering the phenotype as evidenced by the circNFATc3 silencing and overexpression studies (Figure 2, 7) versus results of the linear NFATc3 knockdown. The scrambled siRNA with altered junctional sequence (Figure 2B) and pcDNA3.1 vector without ALU repeats that did not circularize the linear construct failed to show any phenotype in loss-of-function and gain-of-function studies respectively. circNFATc3 plays an essential role in oxidative phosphorylation as evidenced by extracellular flux analysis measuring mitochondrial stress and cell energy phenotype (figure 6,9).

Emerging evidence suggests that circRNAs are involved in complicated functions; such as acting as endogenous RNAs to sponge miRNAs, protein decoys, protein translation, regulating expression of parental genes, modulating alternative splicing, regulating RNA–protein interactions, and acting as scaffolds in the assembly of protein complexes[12,26–29]. Interactome analysis of circNFATc3 interaction with miRNA-circRNA networking using starBase v3.0[30] shows that circNFATc3 is highly associated with Let-7 family members of microRNAs (supplementary Table 2). Let-7 microRNA family exerts its tumor suppressor and antiproliferative activities by repressing several oncogenes; including RAS, and by controlling key regulators of the cell cycle, cell differentiation and apoptotic pathways [31,32]. Let-7 microRNA is known to be involved in a negative feedback loop which downregulates NFAT family gene expression[33]. Similar to its parental gene family, circNFATc3 may also be involved in regulating gene expression *via* let-7-mediated feedback loop. Using StarBase to find RBP-circRNA interactions supported by CLIP-Seq data shows circNFATc3 association with several RBPs (RNA Binding Proteins) (supplementary Table 3). Likewise, using the circRNA intractome[34], (https://circinteractome.nia.nih.gov/) tool shows association between several RBPs and circNFATc3 as well (supplementary Figure 7). As evidenced by circBase and circinteractome, circNFATc3 can potentially sponge microRNAs and RBPs making it important in regulation in multiple biological activities; including cell proliferation, motility, apoptosis, senescence and cell responses to oxidative stress via posttranscriptional regulation such as RNA alternative splicing, conservation, transport and translation[35,36].

To conclude, circNFATc3 is one of the uncharacterized circular RNA which holds immense potential to be a future therapeutic agent. circNFATc3 is involved in regulating cell proliferation, cancer cell invasion, migration and oxidative phosphorylation, which highlights its important role in cancer progression.

## Acknowledgement

The work was supported by grants from Basic Medical Research Program (BMRP) grant from Qatar Foundation to WCM-Q.. We thank Dr. Anna Halama from Dr. Karsten Suhre’s Bioinformatics Core lab at WCM-Q for providing the LL24, NCI-H226, T47D, HCC2935 cells..

## Funding

This study was supported by Weill Cornell Medicine and Qatar Foundation (BMRP 1 - Malek Pilot FY17). The funders had no role in the design of the study, data analysis, interpretation of data and writing the manuscript.

## Conflict of Interst

The authors declare there is no competimg interst regarding this manuscript.

