## Supplementary material for "Regulation of circular RNA circNFATc3 in cancer cells alters proliferation, migration and oxidative phosphorylation": files: supplemetary figures and tables.pptx

### Slide 1
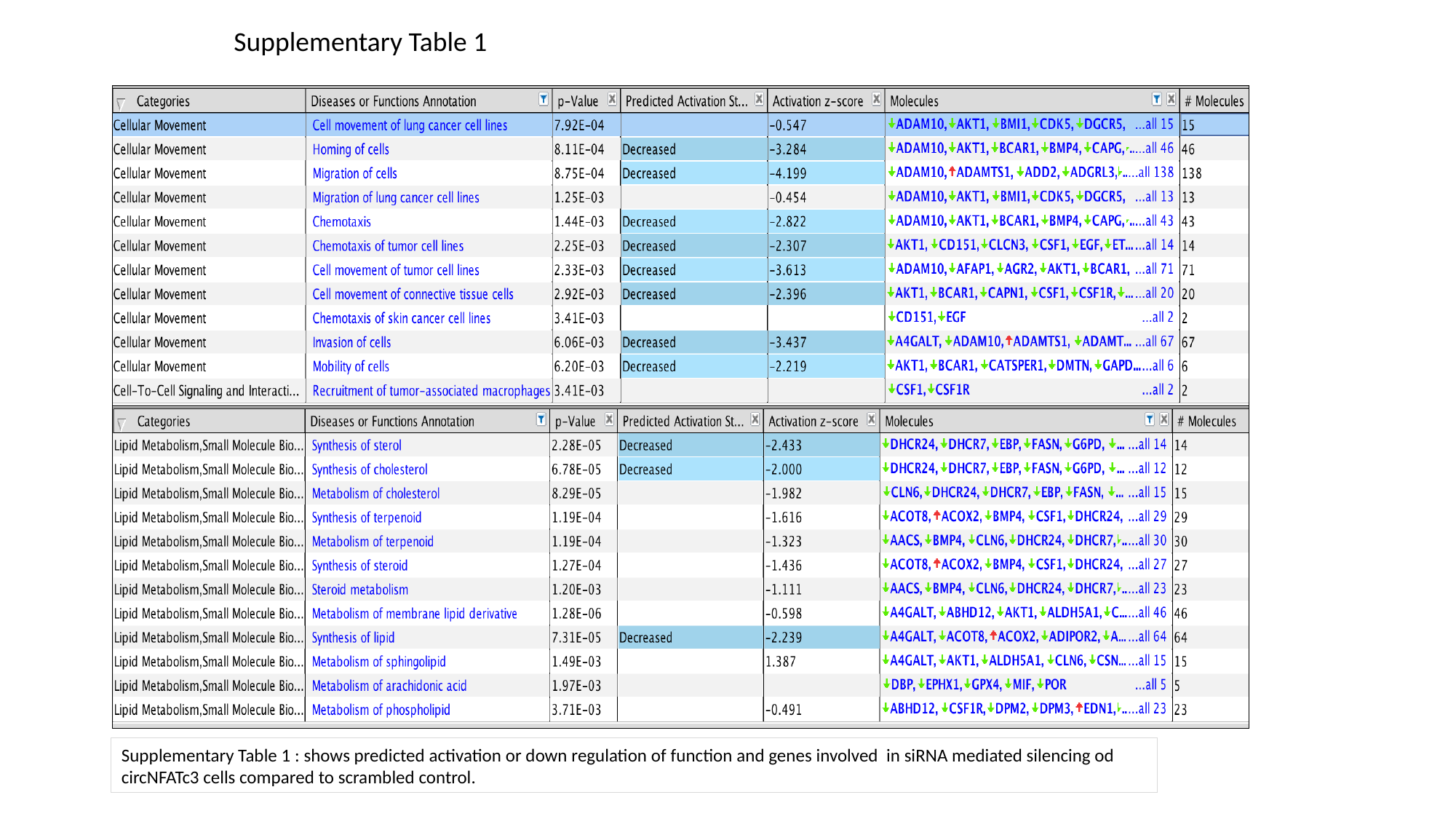

Supplementary Table 1
Supplementary Table 1 : shows predicted activation or down regulation of function and genes involved in siRNA mediated silencing od circNFATc3 cells compared to scrambled control.

### Slide 2
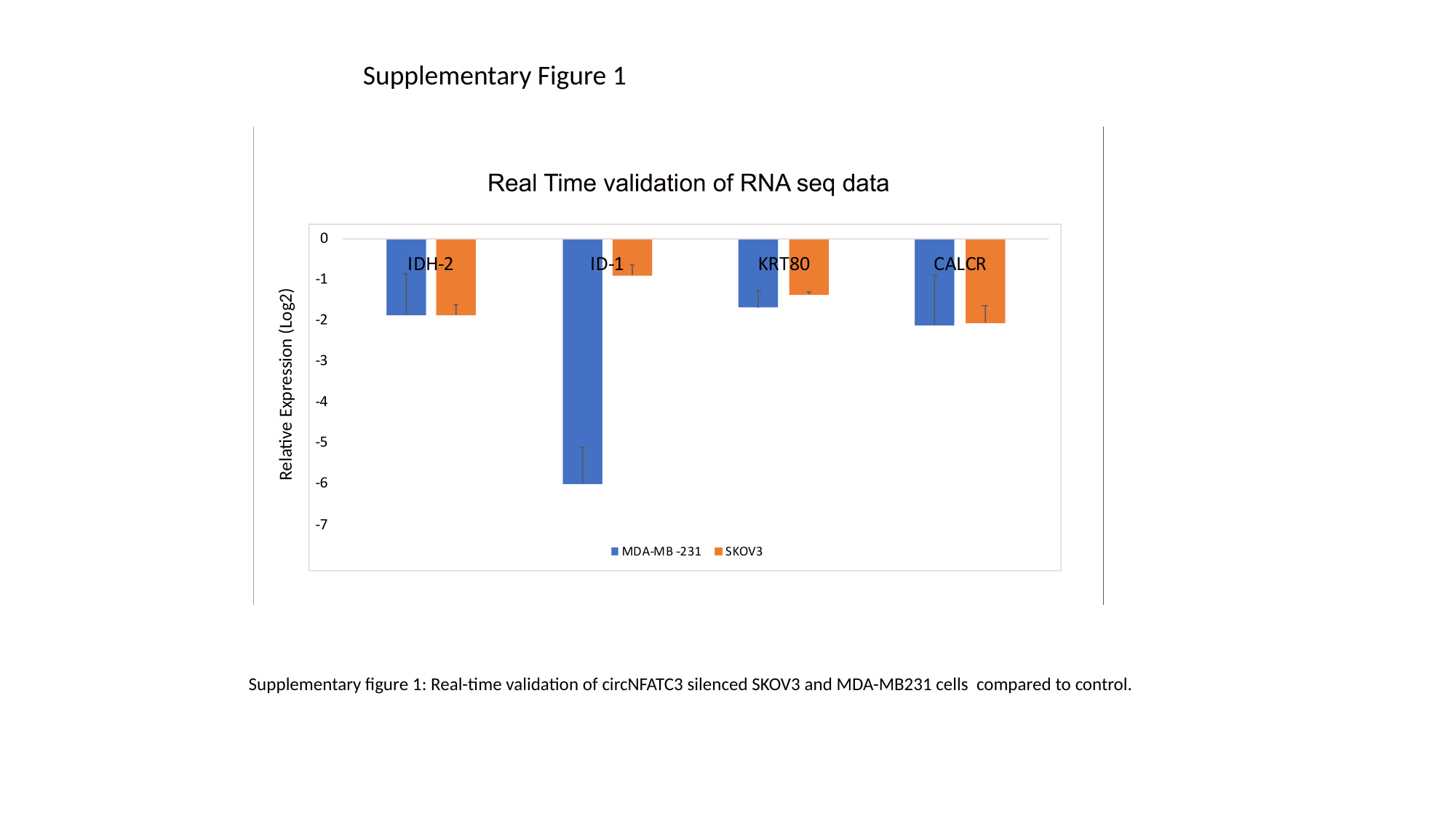

Supplementary Figure 1
Relative Expression (Log2)
Supplementary figure 1: Real-time validation of circNFATC3 silenced SKOV3 and MDA-MB231 cells compared to control.

### Slide 3
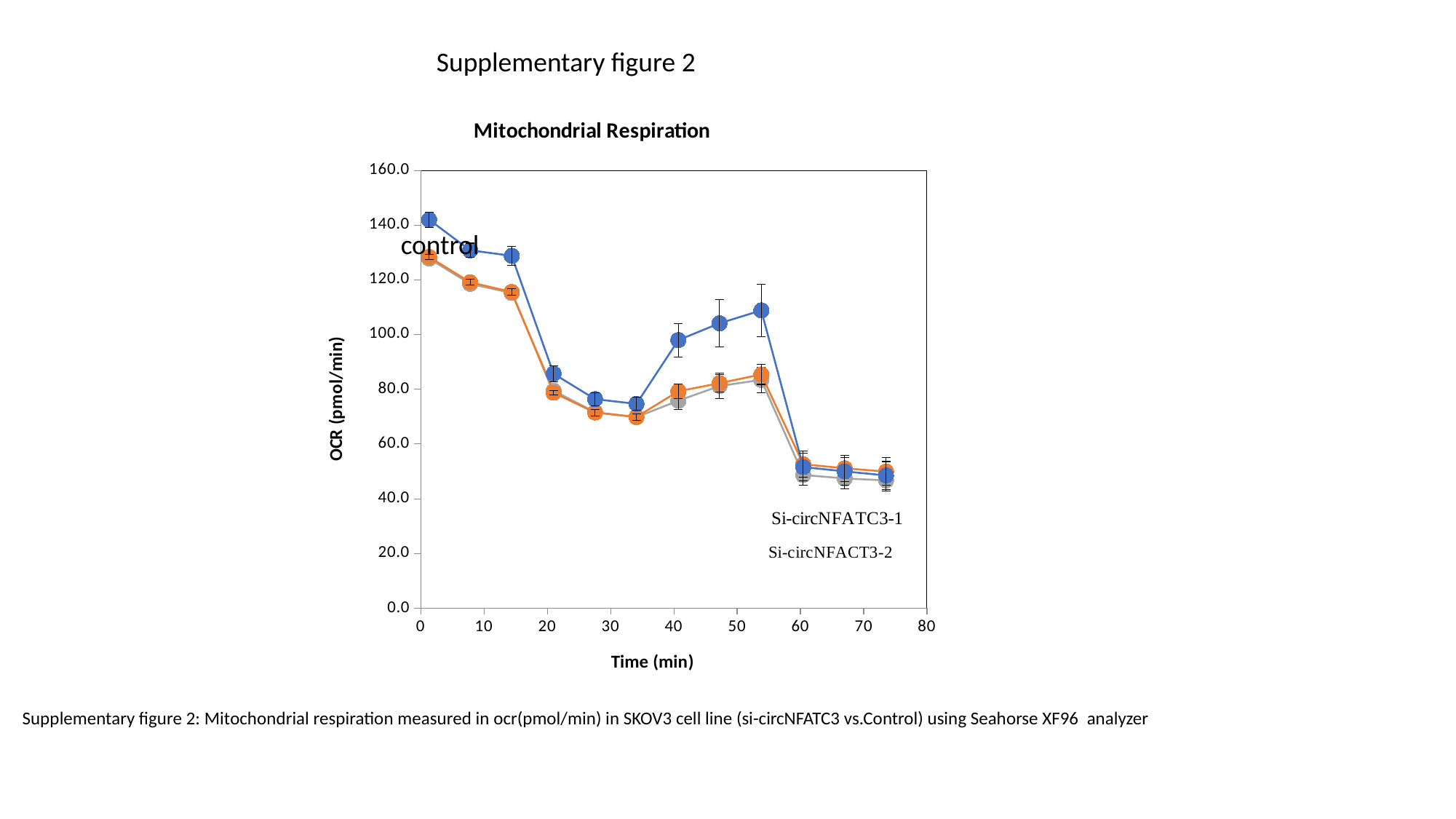

Supplementary figure 2
###### Chart: Mitochondrial Respiration
| Category | Group 1 | Group 2 | Group 3 | Unselected |
|---|---|---|---|---|control
Supplementary figure 2: Mitochondrial respiration measured in ocr(pmol/min) in SKOV3 cell line (si-circNFATC3 vs.Control) using Seahorse XF96 analyzer

### Slide 4
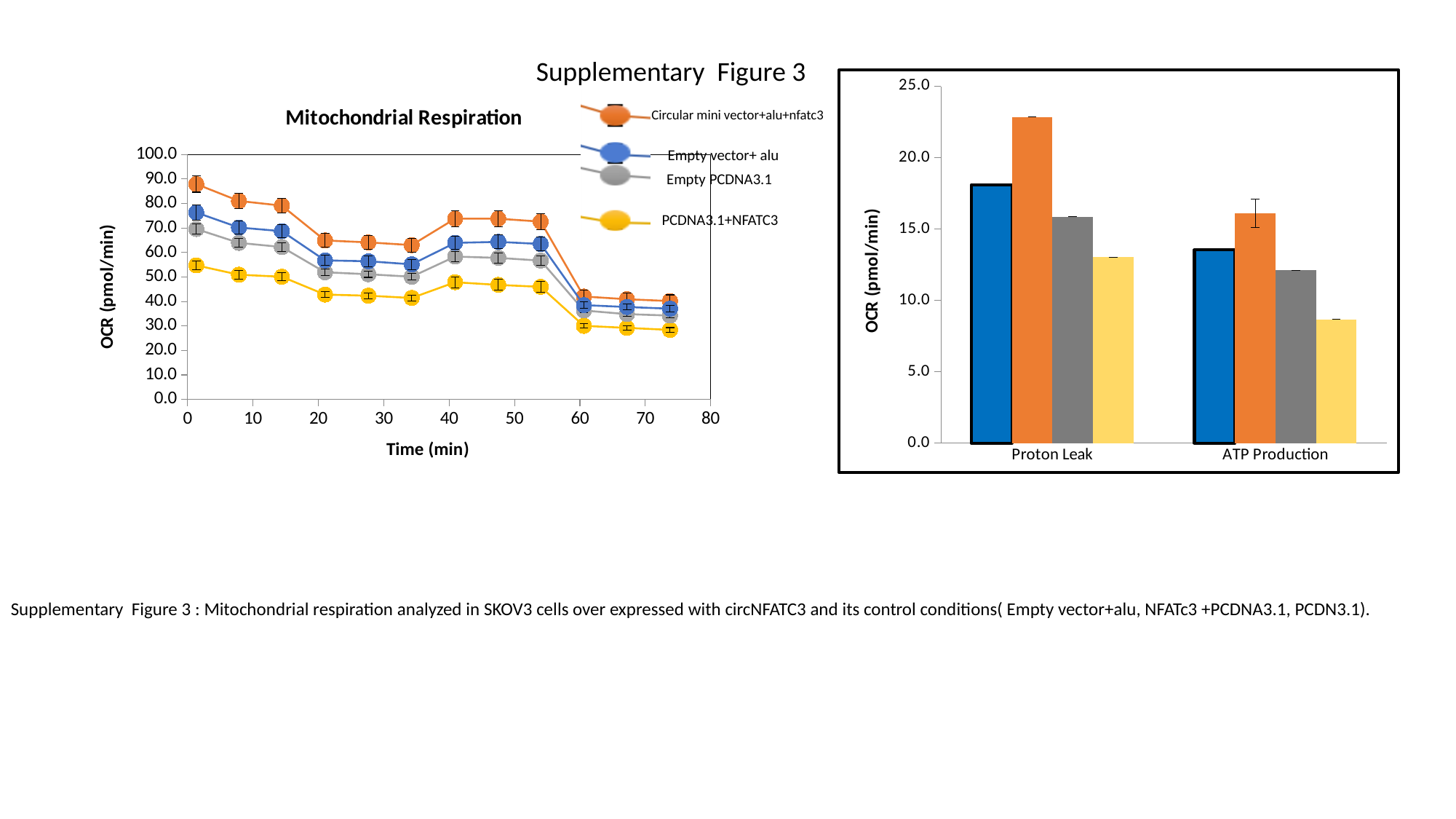

Supplementary Figure 3
###### Chart
| Category | Group 1 | Group 2 | Group 3 | Group 4 |
|---|---|---|---|---|
| Proton Leak | 18.101961135864258 | 22.848114013671875 | 15.853611946105957 | 13.026777267456055 |
| ATP Production | 13.542154312133789 | 16.108840942382812 | 12.119064331054688 | 8.678299903869629 |
###### Chart: Mitochondrial Respiration
| Category | Group 1 | Group 2 | Group 3 | Group 4 | Group 5 | Unselected |
|---|---|---|---|---|---|---|Circular mini vector+alu+nfatc3
Empty vector+ alu
Empty PCDNA3.1
PCDNA3.1+NFATC3
Supplementary Figure 3 : Mitochondrial respiration analyzed in SKOV3 cells over expressed with circNFATC3 and its control conditions( Empty vector+alu, NFATc3 +PCDNA3.1, PCDN3.1).

### Slide 5
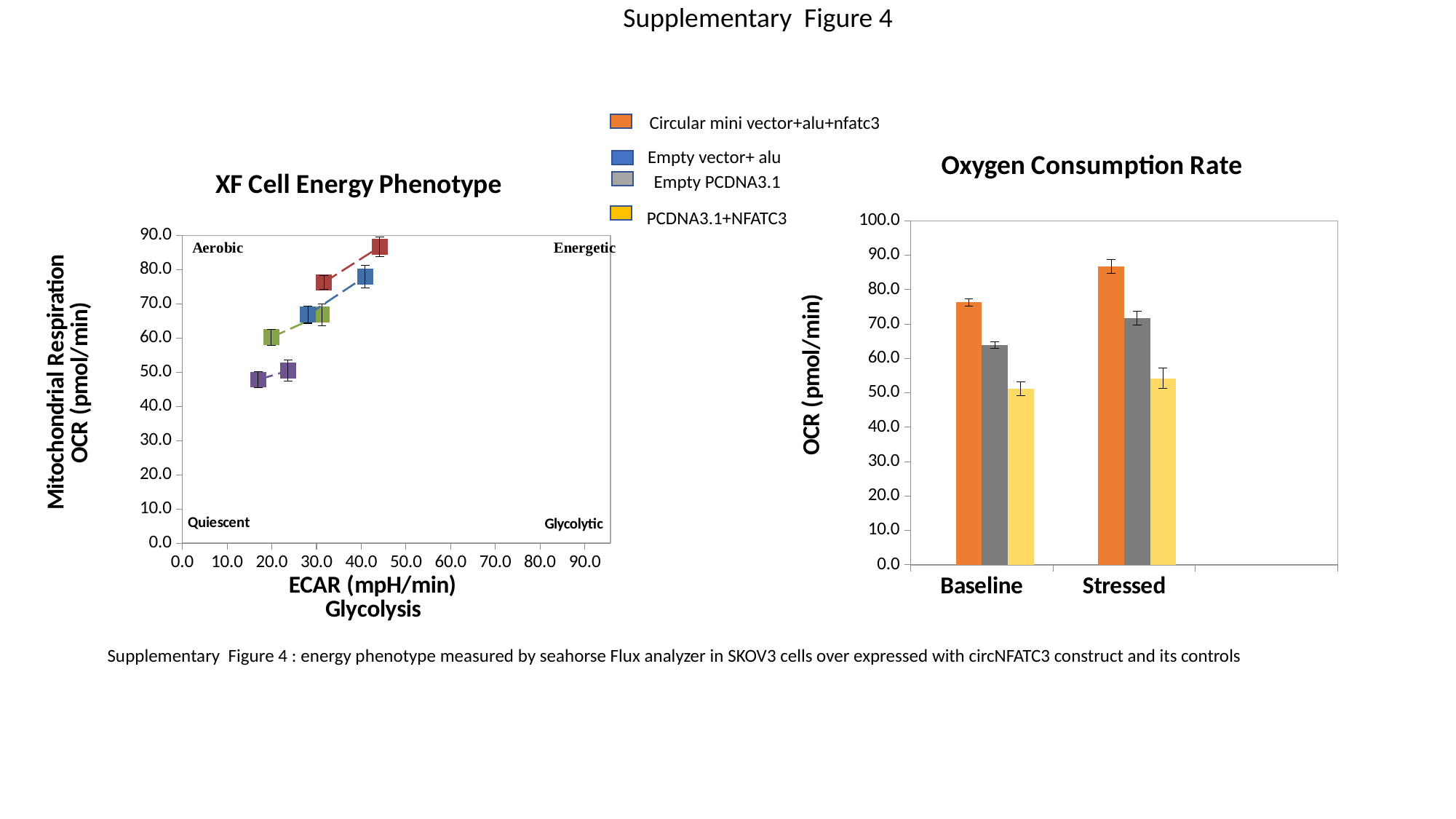

Supplementary Figure 4
Circular mini vector+alu+nfatc3
###### Chart: Oxygen Consumption Rate
| Category | Group 1 | Group 2 | Group 3 | Group 4 |
|---|---|---|---|---|
| Baseline | 64.5655288696289 | 76.29403686523438 | 63.912593841552734 | 51.145511627197266 |
| Stressed | 75.26549530029297 | 86.67937469482422 | 71.7542953491211 | 54.249671936035156 |Empty vector+ alu
###### Chart: XF Cell Energy Phenotype
| Category | | | | | |
|---|---|---|---|---|---|Empty PCDNA3.1
PCDNA3.1+NFATC3
Supplementary Figure 4 : energy phenotype measured by seahorse Flux analyzer in SKOV3 cells over expressed with circNFATC3 construct and its controls

### Slide 6
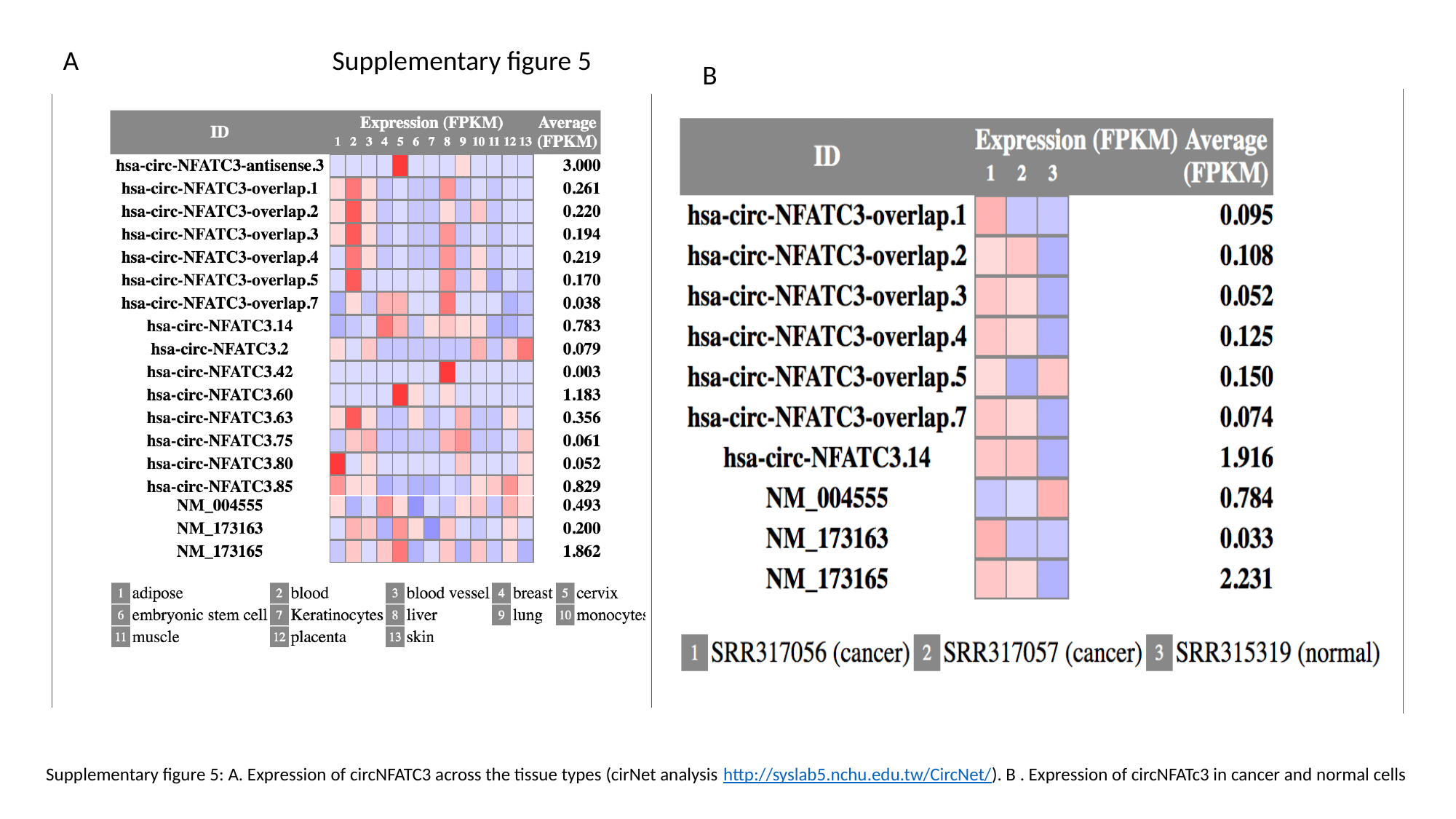

Supplementary figure 5
A
B
Supplementary figure 5: A. Expression of circNFATC3 across the tissue types (cirNet analysis http://syslab5.nchu.edu.tw/CircNet/). B . Expression of circNFATc3 in cancer and normal cells

### Slide 7
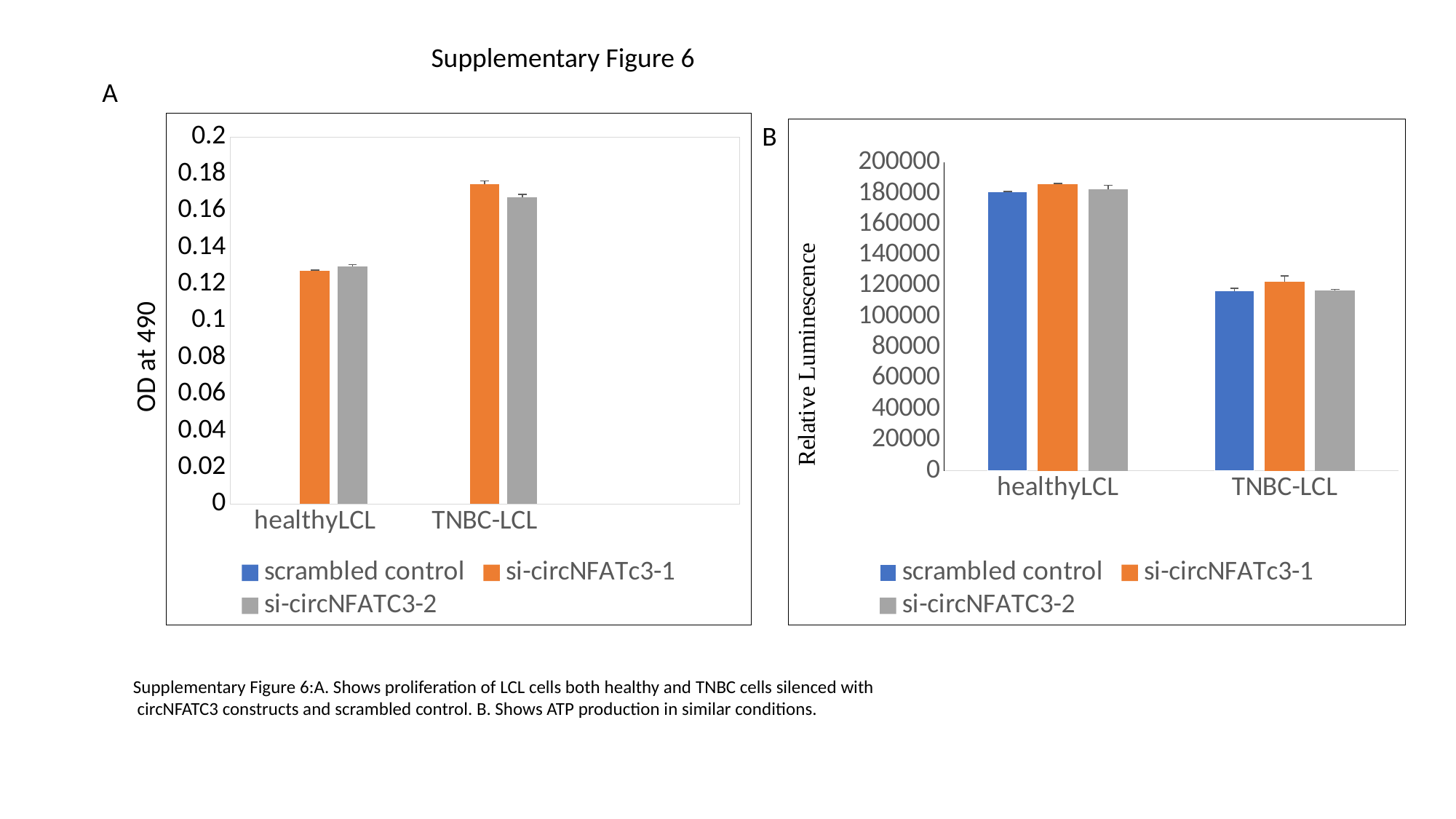

Supplementary Figure 6
A
###### Chart
| Category | scrambled control | si-circNFATc3-1 | si-circNFATC3-2 |
|---|---|---|---|
| healthyLCL | 0.12574999999999997 | 0.12728125 | 0.12965625 |
| TNBC-LCL | 0.17678125 | 0.1745 | 0.16724999999999998 |B
###### Chart
| Category | scrambled control | si-circNFATc3-1 | si-circNFATC3-2 |
|---|---|---|---|
| healthyLCL | 181013.25 | 185888.5 | 182344.9696969697 |
| TNBC-LCL | 116487.20833333334 | 122745.76190476191 | 116765.28571428571 |OD at 490
Supplementary Figure 6:A. Shows proliferation of LCL cells both healthy and TNBC cells silenced with
 circNFATC3 constructs and scrambled control. B. Shows ATP production in similar conditions.

### Slide 8
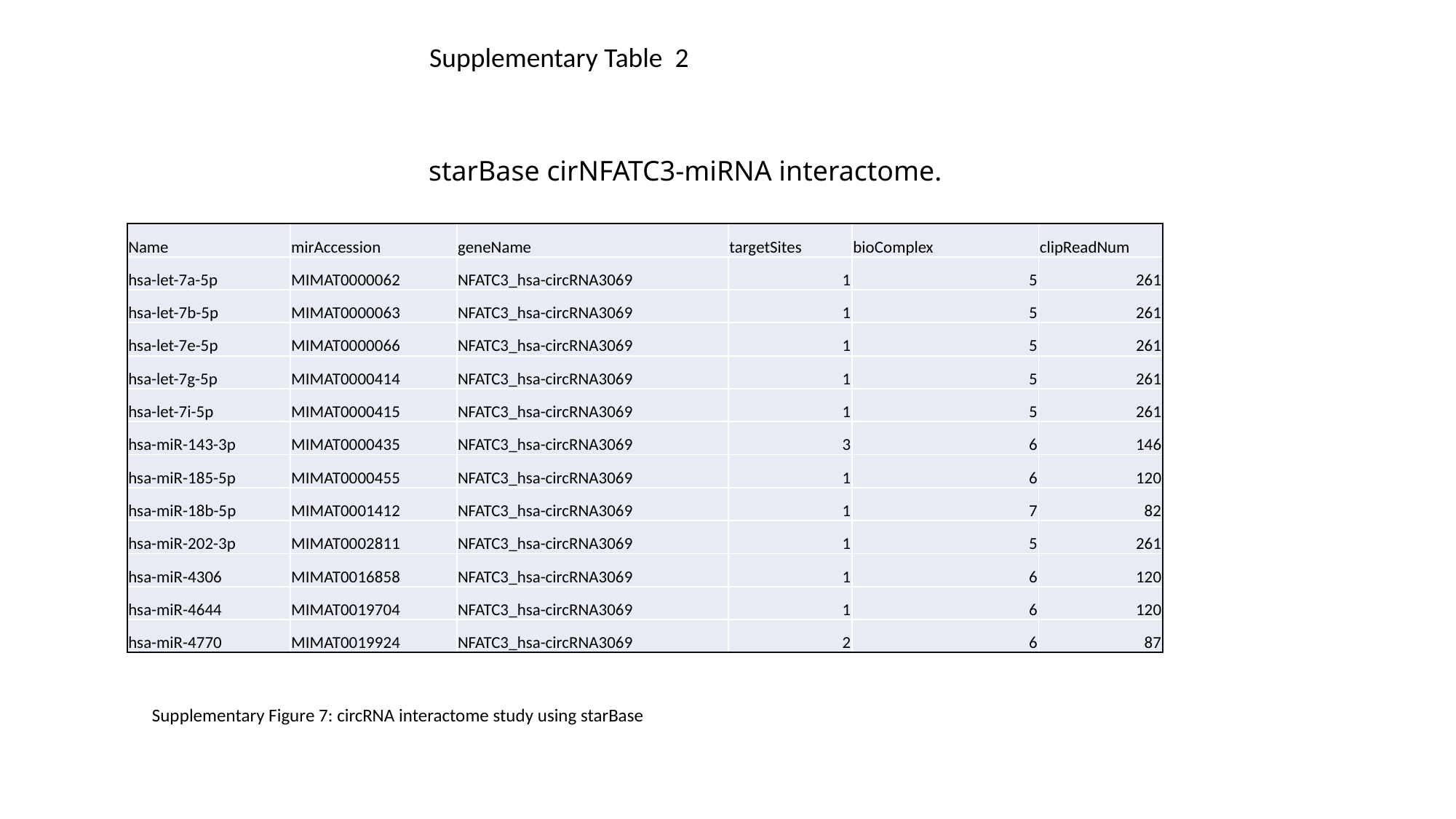

Supplementary Table 2
#### starBase cirNFATC3-miRNA interactome.
| Name | mirAccession | geneName | targetSites | bioComplex | clipReadNum |
| --- | --- | --- | --- | --- | --- |
| hsa-let-7a-5p | MIMAT0000062 | NFATC3\_hsa-circRNA3069 | 1 | 5 | 261 |
| hsa-let-7b-5p | MIMAT0000063 | NFATC3\_hsa-circRNA3069 | 1 | 5 | 261 |
| hsa-let-7e-5p | MIMAT0000066 | NFATC3\_hsa-circRNA3069 | 1 | 5 | 261 |
| hsa-let-7g-5p | MIMAT0000414 | NFATC3\_hsa-circRNA3069 | 1 | 5 | 261 |
| hsa-let-7i-5p | MIMAT0000415 | NFATC3\_hsa-circRNA3069 | 1 | 5 | 261 |
| hsa-miR-143-3p | MIMAT0000435 | NFATC3\_hsa-circRNA3069 | 3 | 6 | 146 |
| hsa-miR-185-5p | MIMAT0000455 | NFATC3\_hsa-circRNA3069 | 1 | 6 | 120 |
| hsa-miR-18b-5p | MIMAT0001412 | NFATC3\_hsa-circRNA3069 | 1 | 7 | 82 |
| hsa-miR-202-3p | MIMAT0002811 | NFATC3\_hsa-circRNA3069 | 1 | 5 | 261 |
| hsa-miR-4306 | MIMAT0016858 | NFATC3\_hsa-circRNA3069 | 1 | 6 | 120 |
| hsa-miR-4644 | MIMAT0019704 | NFATC3\_hsa-circRNA3069 | 1 | 6 | 120 |
| hsa-miR-4770 | MIMAT0019924 | NFATC3\_hsa-circRNA3069 | 2 | 6 | 87 |
Supplementary Figure 7: circRNA interactome study using starBase

### Slide 9
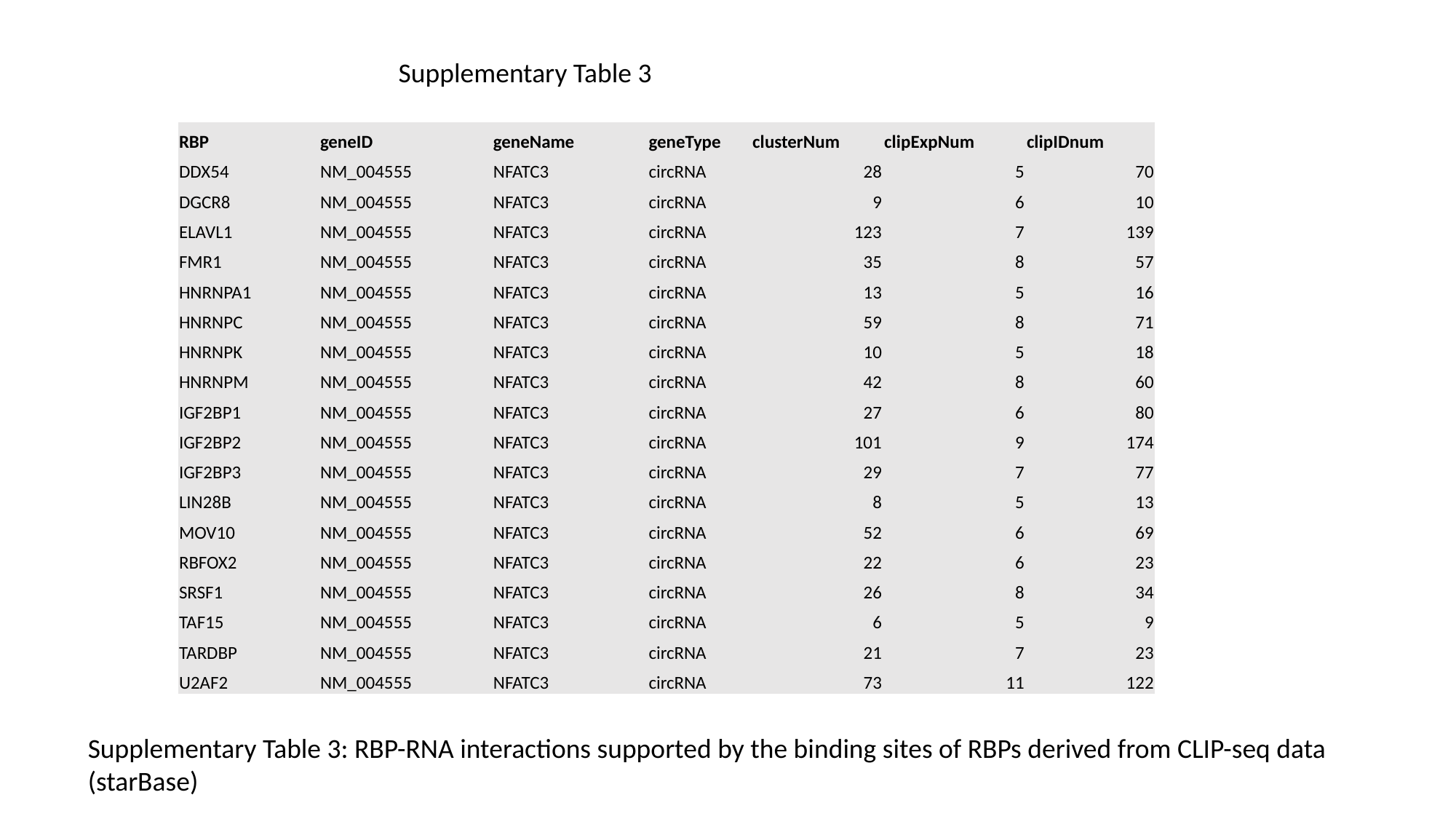

Supplementary Table 3
| RBP | geneID | geneName | geneType | clusterNum | clipExpNum | clipIDnum |
| --- | --- | --- | --- | --- | --- | --- |
| DDX54 | NM\_004555 | NFATC3 | circRNA | 28 | 5 | 70 |
| DGCR8 | NM\_004555 | NFATC3 | circRNA | 9 | 6 | 10 |
| ELAVL1 | NM\_004555 | NFATC3 | circRNA | 123 | 7 | 139 |
| FMR1 | NM\_004555 | NFATC3 | circRNA | 35 | 8 | 57 |
| HNRNPA1 | NM\_004555 | NFATC3 | circRNA | 13 | 5 | 16 |
| HNRNPC | NM\_004555 | NFATC3 | circRNA | 59 | 8 | 71 |
| HNRNPK | NM\_004555 | NFATC3 | circRNA | 10 | 5 | 18 |
| HNRNPM | NM\_004555 | NFATC3 | circRNA | 42 | 8 | 60 |
| IGF2BP1 | NM\_004555 | NFATC3 | circRNA | 27 | 6 | 80 |
| IGF2BP2 | NM\_004555 | NFATC3 | circRNA | 101 | 9 | 174 |
| IGF2BP3 | NM\_004555 | NFATC3 | circRNA | 29 | 7 | 77 |
| LIN28B | NM\_004555 | NFATC3 | circRNA | 8 | 5 | 13 |
| MOV10 | NM\_004555 | NFATC3 | circRNA | 52 | 6 | 69 |
| RBFOX2 | NM\_004555 | NFATC3 | circRNA | 22 | 6 | 23 |
| SRSF1 | NM\_004555 | NFATC3 | circRNA | 26 | 8 | 34 |
| TAF15 | NM\_004555 | NFATC3 | circRNA | 6 | 5 | 9 |
| TARDBP | NM\_004555 | NFATC3 | circRNA | 21 | 7 | 23 |
| U2AF2 | NM\_004555 | NFATC3 | circRNA | 73 | 11 | 122 |
Supplementary Table 3: RBP-RNA interactions supported by the binding sites of RBPs derived from CLIP-seq data (starBase)

### Slide 10
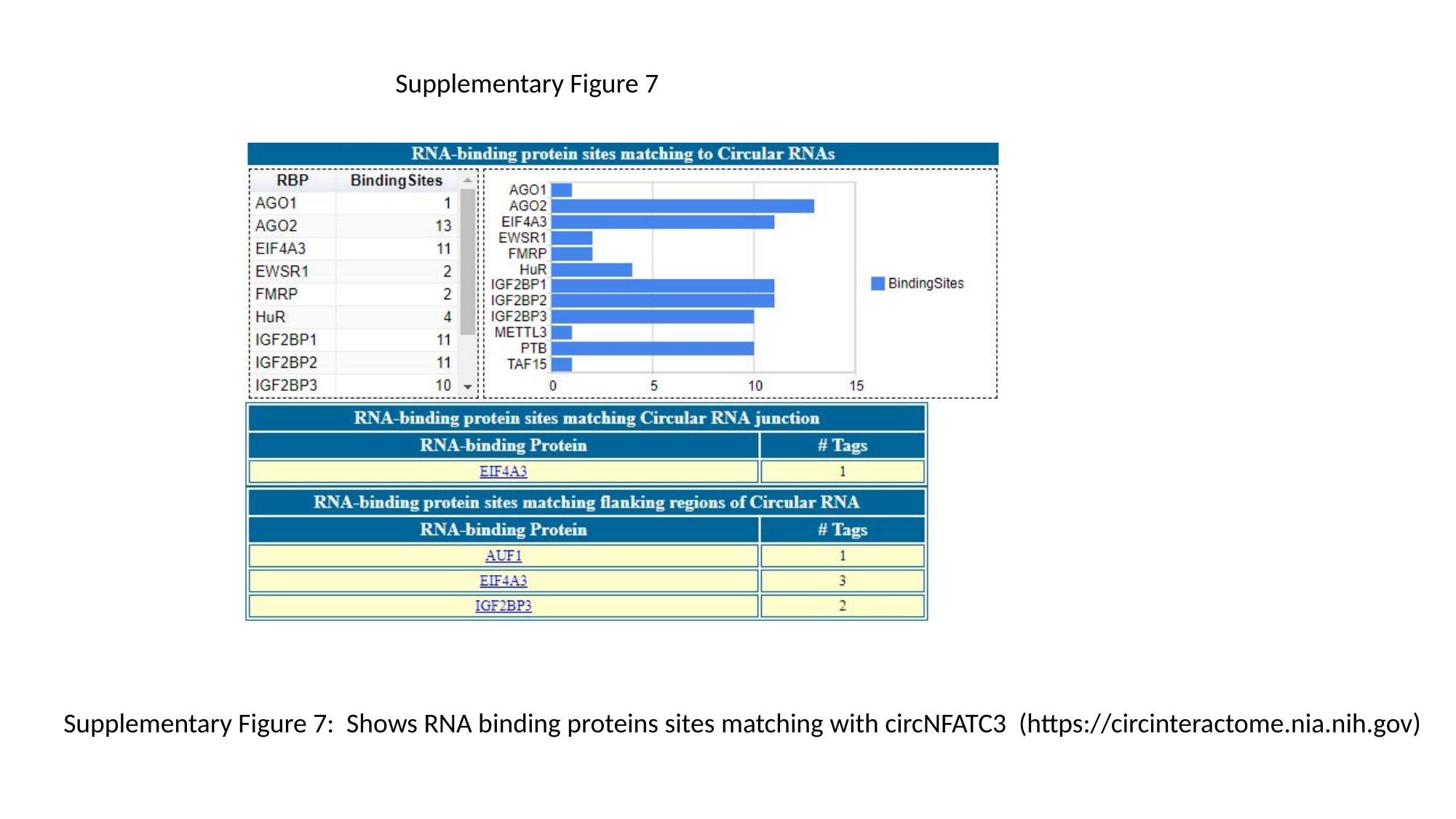

Supplementary Figure 7
Supplementary Figure 7: Shows RNA binding proteins sites matching with circNFATC3 (https://circinteractome.nia.nih.gov)
